# Root-associated *Streptomyces* produce galbonolides to modulate plant immunity and promote rhizosphere colonisation

**DOI:** 10.1101/2024.01.20.576418

**Authors:** Clément Nicolle, Damien Gayrard, Alba Noël, Marion Hortala, Aurélien Amiel, Sabine Grat, Aurélie Le Ru, Guillaume Marti, Jean-Luc Pernodet, Sylvie Lautru, Bernard Dumas, Thomas Rey

**Affiliations:** Laboratoire de Recherche en Sciences Végétales, Université de Toulouse, CNRS, Université Toulouse III, Toulouse INP, 24 Chemin de Borde Rouge, Auzeville, 31320, Auzeville-Tolosane, France; DE SANGOSSE, Bonnel, 47480, Pont-Du-Casse, France; Université Paris-Saclay, CEA, CNRS, Institute for Integrative Biology of the Cell (I2BC), 91198, Gif-sur-Yvette, France; Plateforme d’Imagerie FRAIB-TRI, Université de Toulouse, CNRS, Auzeville-Tolosane 31320, France; Metatoul-AgromiX Platform, LRSV, Université de Toulouse, CNRS, UPS, Toulouse INP, Toulouse, France; MetaboHUB-MetaToul, National Infrastructure of Metabolomics and Fluxomics, Toulouse, France

**Keywords:** Rhizosphere, Galbonolides, *Streptomyces*, *Arabidopsis*, Camalexin

## Abstract

The rhizosphere, which serves as the primary interface between plant roots and the soil, constitutes an ecological niche for a huge diversity of microbial communities. Currently, there is little knowledge on the nature and the function of the different metabolites released by rhizospheric microbes to facilitate colonization of this highly competitive environment. Here, we demonstrate how the production of galbonolides, a group of polyene macrolides that inhibit plant and fungal Inositol Phosphorylceramide Synthase (IPCS), empowers the rhizospheric *Streptomyces* strain AgN23, to thrive in the rhizosphere by triggering the plant’s defence mechanisms. Metabolomic analysis of AgN23-inoculated *Arabidopsis* roots revealed a strong induction in the production of an indole alkaloid, camalexin, which is a major phytoalexin in *Arabidopsis*. By using a plant mutant compromised in camalexin synthesis, we show that camalexin production is necessary for the successful colonization of the rhizosphere by AgN23. Conversely, hindering galbonolides biosynthesis in AgN23 knock-out mutant resulted in loss of inhibition of IPCS, a deficiency in plant defence activation, notably the production of camalexin, and a strongly reduced development of the mutant bacteria in the rhizosphere. Together, our results identified galbonolides as important metabolites mediating rhizosphere colonisation by *Streptomyces*.

Graphical Abstract
Model summarizing the mode of action of galbonolides in stimulating plant defence to support AgN23 colonization of the rhizosphere. Galbonolides secretion by *Streptomyces* sp. AgN23 trigger Inositol Phosphoceramide Synthase (IPCS) inhibition in *Arabidopsis* root cells (orange arrow). The resulting raise in Ceramide precursors of the IPCS may result in the different defence responses associated to AgN23: Hypersensitive Responses (HR), Salicylic Acid (SA) signalling, nuclear Ca^2+^ influx, defence gene expression and camalexin biosynthesis. This production of camalexin (blue arrow) exert a positive effect on AgN23 growth in the rhizosphere, presumably by restricting the growth of bacterial and fungal competitors sensitive to this phytoalexin. In addition, galbonolides secretion in the rhizosphere may also directly interfere with fungal competitors of AgN23.

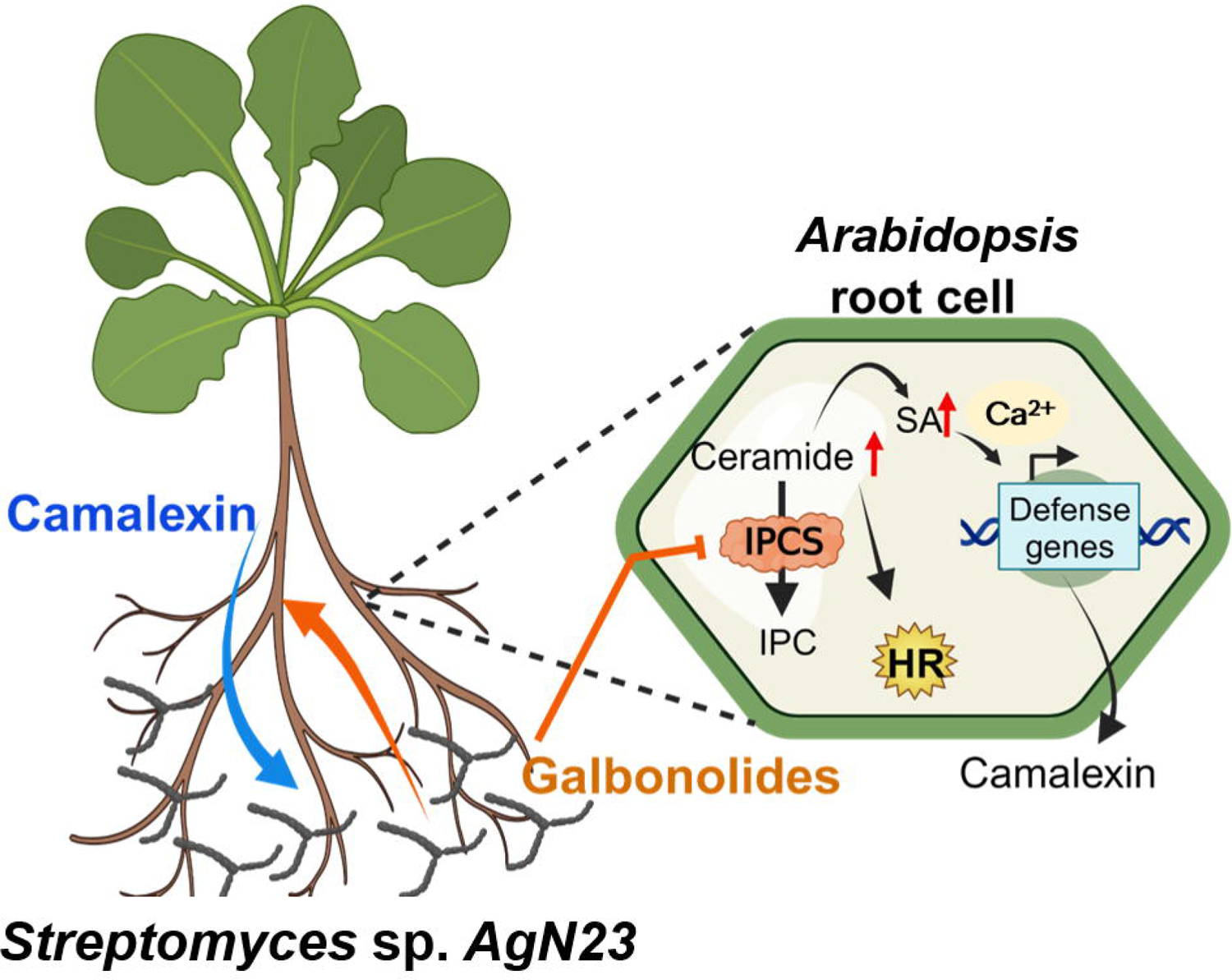

## INTRODUCTION

Cross-kingdom communications play a significant role in shaping interactions between organisms within diverse ecological niches (Mendes and Raaijmakers 2015). Microbe-microbe communication is often mediated by the secretion of small and diffusible specialised metabolites (Andrić et al 2021, Andrić et al 2023, Getzke et al 2023, Krespach et al 2021, Krespach et al 2023). Throughout their lifecycle, eukaryotic organisms such as plants, are known to associate with the abundant and diverse community of microorganisms (Chaparro et al 2014, Durán et al 2018, Fitzpatrick et al 2018). However, there is currently limited knowledge on how plants establish communication with microorganisms and regulate their populations in and around their tissues (Fitzpatrick et al 2020). It has been noted that plants, even when grown in geographically distant soils, tend to assemble a core microbiota comprising bacteria, fungi and oomycetes, suggesting the existence of broad trans-kingdom communication mechanisms within plant-microbe interactions (Thiergart et al 2020). In this context, it becomes primordial to understand the molecular basis of plant-microbiota assembly to achieve the intelligent engineering of crops microbiota (Russ et al 2023). Such approach would be an important milestone towards sustainable agricultural practices in nutrition and protection against pathogens and abiotic stresses (Berendsen et al 2018, Busby et al 2017). An example of this approach is the recently reported study on how a *Streptomyces* strain alleviates abiotic stress in a plant by producing pteridic acid (Yang et al 2023).

The *Streptomyces* genus belongs to Actinomycetes, a family of filamentous sporulating Gram+ bacteria which constitutes the second most prominent component of root microbiota after proteobacteria (Bulgarelli et al 2012, Lebeis et al 2015, Ling et al 2022). Among Actinomycetes, *Streptomyces* spp. are enriched in endophytic and epiphytic root compartments and represent up to 30% of the total bacterial Operational Taxonomic Units (OTUs)(Lundberg et al 2012). Enrichment of *Streptomyces* spp. in soil and rhizosphere correlates with resistance to drought and pathogen attack (Cha et al 2016, Fitzpatrick et al 2018). Furthermore, streptomycetes are hallmark producers of antimicrobial specialised metabolites involved in protection against plant pathogens (Avalos et al 2020, Cordovez et al 2015, Kim et al 2019). *Streptomyces* spp. have also been demonstrated to elicit Salicylic Acid (SA) and ISR (Induced Systemic Resistance) dependent responses leading to the activation of plant defence metabolism (Conn et al 2008, Kurth et al 2014). These important attributes have stimulated great interest in the use of streptomycetes for crop protection (Rey and Dumas 2017, Viaene et al 2016).

In a previous paper, we reported the screening of a collection of 35 *Streptomyces* strains isolated from agricultural soils for their plant defence elicitation (Vergnes et al 2020). Among these, the AgN23 strain has been reported to display a remarkable potential to elicit *Arabidopsis* defences associated to salicylate, jasmonate and ethylene signalling (Vergnes et al 2020). Foliar inoculation with the bacteria resulted in the formation of *SALICYLIC INDUCTION DEFICIENT 2 (SID2)* dependent necrotic symptoms in *Arabidopsis* and protection against *Alternaria brassicicola* colonisation (Vergnes et al 2020). A recent analysis of the AgN23 genome showed that the strain belongs to the clade *S. violaceusniger* (Gayrard et al 2023). The AgN23 genome harbours large gene families associated to rhizosphere colonization, such as biosynthetic gene clusters (BGCs) involved in the synthesis of plant bioactive and antimicrobial compounds, plant cell wall degrading enzymes, and phytohormone synthesis.

In this work, we investigate the molecular basis of AgN23 interaction with plant roots by characterising rhizosphere colonisation by the bacteria and the resulting plant responses. We find that AgN23 triggered plant biosynthesis of the antimicrobial camalexin and show that this phytoalexin is an important feature for rhizosphere colonisation by the *Streptomyces*. In addition, we established that AgN23 produce galbonolides that can interfere with plant sphingolipid metabolism by targeting the Inositol Phosphorylceramide Synthase. Finally, we show that galbonolides biosynthesis by AgN23 is instrumental for plant defence stimulation, including camalexin production and rhizosphere colonisation by the bacterium.

## MATERIAL AND METHODS

### Plant material, growth conditions and phenotyping

Seeds of *Arabidopsis thaliana* accession Col-0 (N1092) were obtained from the Nottingham Arabidopsis stock center (NASC) and mutant *pad3-1* (N3805) were kindly provided by Dr. Pawel Bednarek. *Arabidopsis* plants grown in potting soil (PROVEEN; Bas Van Buuren B.V., Holland) were cultivated in a growth chamber under 16 hours photoperiod and 23 °C unless otherwise indicated. Similar conditions were applied to the cultivation of *Nicotiana benthamiana*. One-month-old *N. benthamiana* leaves were syringe-infiltered with bacterial CMEs. Cell death areas were photographed 24–48 hours after infiltration, with an Expression 11000 XL scanner (Epson) at 300 dots/inch.

To perform soil inoculation assays with AgN23, 70 g of potting soil inoculated with AgN23 spore inoculum at 10^4^ CFU/g was distributed in pots placed in small plastic bags to avoid cross-contamination during watering. 5 to 10 *Arabidopsis* seeds were sown per pot and the pots were placed in a growth phytotronic chamber. A single seedling was kept per pot 5 days after germination. Pots were watered weekly with 10 mL of tap water. The watering pots were photographed to monitor the aerial part phenotype. Green area was measured with ImageJ (v. 1.51k) at 4, 6 or 7 weeks after inoculation. Details regarding *in vitro* cultivation of *Arabidopsis* are in Supplementary methods.

### AgN23 cultivation and transgenesis to obtain reporter lines and Δ*gbnB* mutants

AgN23 was grown in Bennett medium for the purpose of liquid state cultivation and of Culture Media Extract (CME) production (D-Glucose 10 g/L; Soybean peptones 2.5 g/L; Yeast Extract 1.5 g/L; Sigma). The culture was set in 250 mL Erlenmeyer flasks by inoculating 50 mL Bennett medium with 100 µL of fresh spore suspension at 10^5^ CFU/mL at 250 rpm and 28 °C in a shaking incubator at 250 rpm for 7 days. For the purpose of spore production and genetic manipulations AgN23 strain was cultivated on the solid medium Soya Flour Mannitol (SFM) medium (D-Mannitol (Sigma) 20 g/L; organic soya flour (Priméal) 20 g/L; BactoTM Agar (Difco Laboratories) 20 g/L).

*Escherichia coli* strains were grown in LB with appropriate antibiotics as necessary. *E. coli* transformation and *E. coli* / *Streptomyces* conjugation were performed according to standard procedures (Kieser et al 2000, Sambrook et al 2000). Phusion High-fidelity DNA Polymerase (Thermo Fisher Scientific) was used to amplify DNA fragment except for PCR verification of plasmids or strains for which Taq polymerase (Qiagen) was used. DNA fragments and PCR products were purified using the Nucleospin Gel and PCR clean up kit (Macherey-Nagel).

For pOSV700 plasmid construction, a 0.4 kb DNA fragment encompassing the ermEp* promoter and the tipA ribosome binding site (RBS) was amplified from pOSV666 using the primers JWseq6 and JWseq7. The fragment was digested by EcoRV and cloned into EcoRV-digested pSET152, resulting in pOSV700. The sequence of the insert was verified.

For GFP and mCherry transgenesis, the sequences of the soluble-modified GFP (smGFP) and mCHERRY genes were optimized for expression in *Streptomyces*, synthesized as gblocks (IDT) and cloned into pGEM-T easy, resulting in pmsolGFP and pmCHERRY, respectively. The smGFP and mCHERRY genes were amplified from these plasmids using the primer pairs onSC001/onSC011 and onSC005/onSC013, respectively. PCR amplicons were digested by NdeI and PacI and cloned into NdeI/PacI-digested pOSV700. The resulting plasmids were verified by restriction digestion and sequencing, and named pSC001 (smGFP) and pSC003 (mCHERRY). These were subsequently introduced in *E. coli* ET12567/pUZ8002 and transferred into *Streptomyces* sp. AgN23 by intergeneric conjugation. Conjugants were selected on apramycin 50 µg/ml. The resulting strains were verified by PCR on the extracted genomic DNA using the pSET152-F and pSET152-R primers.

For production of galbonolides knock-out mutants, a 5 kb internal fragment of *gbnB* coding for the structural PKS gene of the galbonolides biosynthetic gene cluster was replaced by a kanamycin resistance cassette. For this purpose, a 2kb fragment (upstream fragment) encompassing the beginning of *gbnB* was amplified by PCR with the onSC007/onSC008 primer pair and cloned into pGEM-T Easy, yielding pSC008. Similarly, a 2kb fragment (downstream fragment) encompassing the end of *gbnB* was amplified by PCR with the onSC009/onSC010 primer pair and cloned into pGEM-T Easy, yielding pSC009. The pSC008 and pSC009 plasmids were digested by EcoRI/EcoRV and DraI/HindIII, respectively, and the 2 kb fragments (upstream and downstream fragments respectively) were purified on agarose gel. The kanamycin resistance cassette was obtained by digesting pOSV514 by EcoRV. The three fragments (upstream, downstream and resistance cassette) were next ligated into EcoRI/HindIII-digested pOJ260. The resulting plasmid, named pSC004, was verified by digestion with BamHI, PstI, EcoRI and EcoRV. Five independent conjugants were verified by PCR using the onSC022/onSC023, onSC021/JWseq16, and onSC030/JWseq17 primer pairs. All oligonucleotides used in this work are listed in Supplementary Table S5.

### Analysis of *Arabidopsis* defence response

Detailed procedures for *Arabidopsis* loss of electrolytes and Calcium signal detection are described in Supplementary Methods. For defence gene expression assays, total RNAs were extracted using the RNeasy Plant Mini Kit (Qiagen) and DNase treated with RQ1 RNase-Free DNase (Promega). For each sample, 1 µg of total RNA was reverse-transcribed with the High Capacity cDNA Reverse Transcription Kit (Applied Biosystems). cDNAs were diluted to 1 ng/ µL and used for qPCR analysis in a 10 µL reaction mix containing 5 µL of LightCycler® 480 SYBR Green I Master mix (Roche), 300 nM of each primer, and 2 µL of the diluted template cDNAs. qPCR was performed in triplicate using a LightCycler ® 480 System (Roche) with preheating at 95 °C for 5 minutes and then 40 cycles of 95 °C for 15 s and 60 °C for 60 s. The Polyubiquitin 10 gene AT4G05320 was retained for normalization. The 2-ΔCp method was used to display gene expression levels 82. Primers used in this study are listed in Supplementary Table S5.

### AgN23 DNA quantification from soil and rhizosphere DNA

To track the development of AgN23, the plants were removed from the soil. The remaining soil from each pot was homogenized and a small amount was sampled and considered as bulk sample. Roots were placed into 50 mL conical sterile polypropylene centrifuge tubes filled with 20 mL 1X phosphate-buffered saline (pH 7.4), and vigorously vortexed to release the adhering rhizospheric soil. Tubes were then centrifuged at 4000 rpm and the washing step was repeated one time. Soil pellets after second centrifugation step were considered as rhizosphere samples. Samples were stored at −80°C until processing. The total microbe DNA from 100 mg of bulk or rhizosphere samples was extracted using the *Quick*-DNATM Fecal/Soil Microbe Miniprep kit (Zymo Research) following manufacturer’s instructions. DNA was eluted in 100 µL DNA Elution Buffer and quantified with DS-11 Spectrophotometer/Fluorometer (DeNovix) and stored at −80°C until processing. The experimental procedures and calculation for AgN23 genome copies quantification in the rhizosphere is detailed in Supplementary methods.

### Preparation of samples for biochemistry studies and mass spectrometry analysis

For root metabolome studies, 10 *Arabidopsis* seedlings from the same MS plate were sampled together in 2 mL microtubes containing two 3 mm-diameter tungsten carbide beads (Qiagen), and flash frozen in liquid nitrogen. For studies of AgN23 culture media extract (CME), the bacterial biomass grown in liquid flask culture was removed from the culture supernatant by centrifugation at 4,200 rpm for 10 min, completely dried in oven at 50 °C, and weighed to assess AgN23 growth. Further details regarding metabolites extraction of AgN23 and *Arabidopsis* are detailed in Supplementary methods.

### Microscopy

For stereo microscopy, we used a Nikon SMZ16 microscope equipped with a camera. Confocal microscopy was performed on a TCS SP8 confocal microscope (Leica, Microsystems, UK). For GFP-tagged AgN23 cells, the excitation wavelength was 488 nm, with emission absorbance between 500 nm and 550 nm, whereas an excitation wavelength of 543 nm was used for mCherry-tagged AgN23 cells proteins, with emission absorbance between 560 nm and 600 nm. Images were acquired with a ×40 or ×20 water immersion lens. All confocal images were analysed and processed using the ImageJ software package (http://rsb.info.nih.gov/ij/; v. 1.51k).

### *Arabidopsis* Inositol Phosphorylceramide Synthase inhibition assay

To study the effect of AgN23 CME on *Arabidopsis* IPCs, we purified microsomal fractions of transgenic yeast expressing AtIPCS2 (AT2G37940) in enzymatic activity assays. First, a preculture of yeast MSY23-3C pESC-LEU_AtIPCS2 strain was performed by picking a single colony and propagating it in 5 mL SGR-TRP-LEU medium (0.1% galactose, 1% raffinose)(Pinneh et al 2019). The preculture was incubated at 30 °C, 200 rpm until the OD_600_ reached 0.8. The preculture was then mixed with 245 mL of fresh SGR medium and incubated at 30 °C, 200 rpm until the OD_600_ reached 0.8. Yeast cells were then harvested by centrifugation, washed with cold phosphate-buffered saline, and stored at −80 °C until microsome preparation. Crude microsomal membranes from yeast MSY23-3C pESC-LEU_AtIPCS2 strain were prepared as previously described with additional CHAPSO washing steps (Mina et al 2010). Total protein quantification was performed by Bradford assay and aliquots at 0.5 mg/mL were made and stocked at −80 °C. Details regarding analytical parameters to study AtIPCS2 enzymatic activity are described in Supplementary methods.

## RESULTS

### AgN23 colonises *Arabidopsis* rhizodermis and rhizosphere

In view of the fact that AgN23 was isolated from grapevine rhizosphere, we looked into the interaction of the strain with roots by inoculating *in vitro* grown *Arabidopsis thaliana* Col-0 seedlings with AgN23 spores. A drop of the spore suspension was deposited at the root tip of young seedlings. A strong development of bacterial microcolonies was observed at the inoculation site 10 days after inoculation (Figure 1a). We generated GFP and RFP-labelled transgenic AgN23 strains and observed the colonization patterns of both strains by epifluorescence microscopy. Results showed that bacteria can spread beyond the initial inoculation spot and colonise other developing sections of the root system, such as lateral roots and apical meristem (Figure 1b). Visual and microscopic inspection of the AgN23-treated plants suggested that the inoculated bacteria did not lead to any characteristic symptoms such as root browning or rhizodermis damages in *Arabidopsis*. Moreover, penetration of AgN23 into the root tissues was not observed. Nevertheless, we observed that AgN23 inoculation did result in a slight (about 10%) reduction in root elongation (Figure 1c).

**Figure 1:**
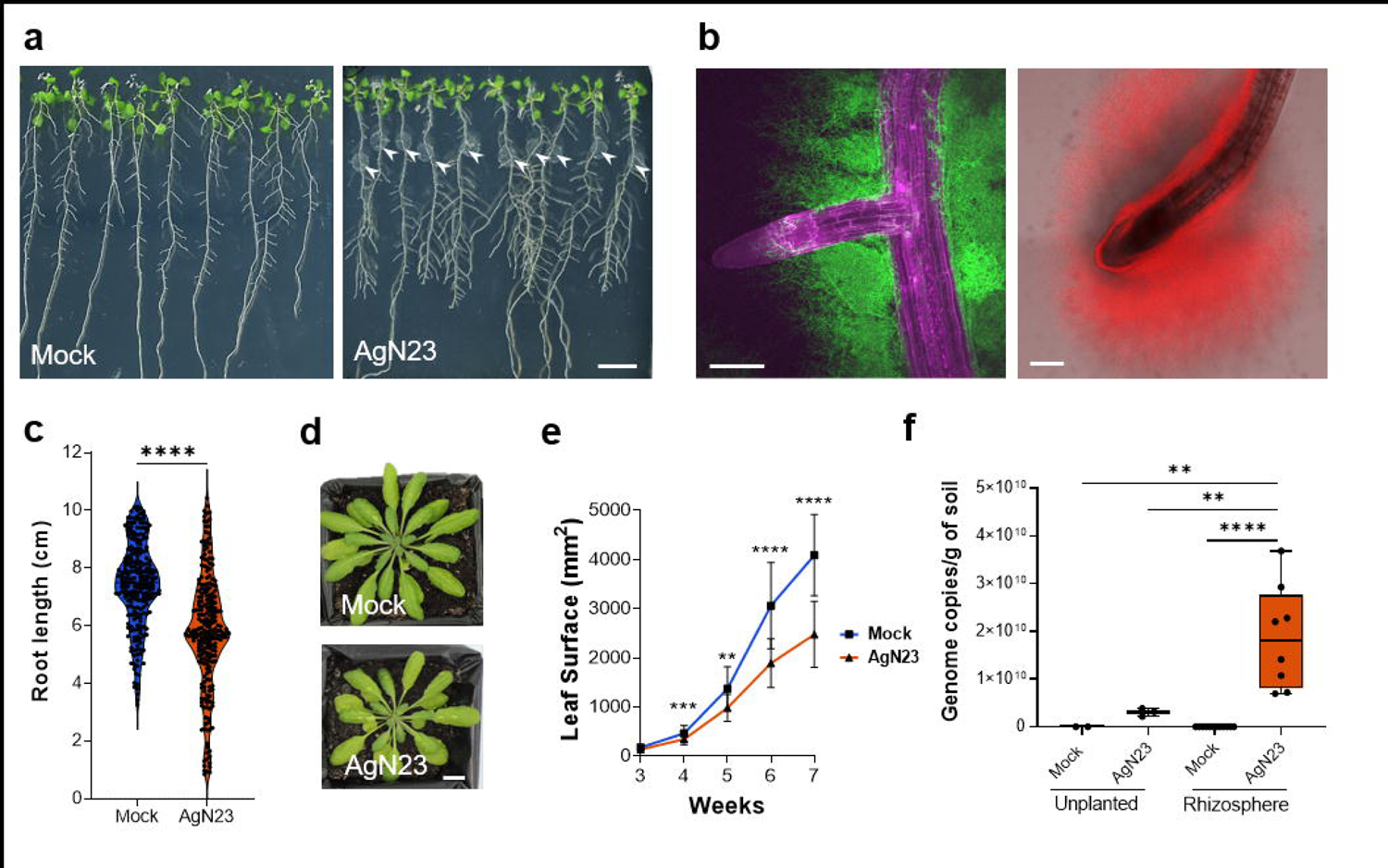
AgN23 colonizes rhizoplane and rhizosphere of *Arabidopsis thaliana* and slightly inhibits plant growth. **a.** Observation of *A. thaliana* Col-0 colonization by AgN23 at 10 days after inoculation with spores at the root apex. Arrowheads indicate AgN23 initial spore inoculation, Scale bar = 1 cm. **b.** Confocal fluorescence images of AgN23-mCherry (red) colonizing root apical meristem and AgN23-GFP (green) developing around lateral root. Scale bars: 250µm **c.** Primary root length of *Arabidopsis* seedlings 10 days after inoculation with AgN23 spores at the root apex from Violin plots created from data from 30 independent assays each involving at least 10 plants per treatment (n = 300). The whiskers encompass the minimum and maximum values, and the midline shows the median. Statistical differences between the treatments were analyzed using Mann–Whitney test and ‘****’ represents significant differences at *P* value < 0.0001. **d.** Typical photographs of 6-week-old Arabidopsis rosettes following growth within non- or inoculated potting soil. Scale bar: 1 cm. **e.** Leaf area measurement of *Arabidopsis* rosettes grown in AgN23-inoculated potting soil. Graphs show the mean ± SD calculated from at least eight biological replicates (n = 8). Statistical comparison between inoculation and mock conditions was performed based on T-test (‘**’ = *P* value < 0.01; ‘***’ = *P* value < 0.001; ‘****’ = *P* value < 0.0001). **f.** AgN23 genome copy number in *Arabidopsis* rhizosphere 4 and 8 weeks after soil inoculation. Box plots were created from data involving at least 8 plants per treatment (n = 8). Whiskers encompass the minimum and maximum values, and the midline shows the median. Statistical differences between the treatments were analyzed using Mann–Whitney test; ‘****’ and ‘**’ represent significant differences at *P* value < 0.0001 and *P* value < 0.05, respectively.

To study the colonization of the rhizosphere by AgN23, we inoculated potting soil with 10^5^ AgN23 spores/g of soil prior to sowing Arabidopsis seeds. Consistent with our previous *in vitro* observation, the presence of AgN23 reduced rosette growth without causing obvious symptoms to the leaves (Figure 1d, 1e) (Vergnes et al, 2020). The development of AgN23 in the inoculated soil was monitored by extracting microbial DNA from both unplanted and rhizosphere soil samples. A quantification of AgN23 genome copies was implemented by amplifying a specific genomic region of the strain from soil samples and from a standard curve of AgN23 purified DNA. Knowing the molecular weight of AgN23 genome, we extrapolated genome copies number from the estimated mass of AgN23 DNA detected in soil. A total of 3.06×10^8^ genome copies of AgN23 were detected in the unplanted soil 7 weeks post inoculation, whereas 1.87×10^10^ genome copies of the bacteria were detected in the rhizosphere. This shows that AgN23 preferentially colonised the *Arabidopsis* rhizosphere rather than the unplanted soil (Figure 1F). Taken together, our data confirm that AgN23 is an epiphytic and rhizospheric bacterium that triggers slight reduction in plant growth, albeit without symptoms.

### Activation of camalexin biosynthesis by AgN23 promotes bacteria settlement in the rhizosphere

In a previous work, we characterised the plant defence stimulating activity of AgN23 and found that the bacterial culture media extract (CME) induced robust transcriptional responses associated with *Arabidopsis* specialized metabolism (Vergnes et al 2020). Detailed analysis showed transcriptional induction of genes coding enzymes involved in camalexin biosynthesis following treatment with AgN23 CME after 1 and 6 hours post-treatment which is a major phytoalexin of *Arabidopsis* belonging to indole alkaloid (Supplementary Figure S1).

To analyse the metabolomic response of root tissues to AgN23, we extracted the metabolites from whole seedlings cultivated *in vitro* in contact with AgN23 for 10 days. The extracts were subjected to a full-scan LC-HRMS metabolomics analysis in ESI+ and ESI-modes and then combined in a single list of variables. A total of 511 variables were retrieved across all the samples out of which, 416 received level 3 annotations according the Metabolomic Standard Initiative based on internally built database and the exact mass and fragmentation profile of the ions (Supplementary Table S1). Unsupervised PCA of the complete variable dataset allowed us to clearly discriminate the control samples from those inoculated with AgN23, with component 1 and component 2 supporting 32.9 % and 21.6% of the variability, respectively (Supplementary Figure S2).

To identify the underlying chemical classes supporting the separation of control and AgN23 inoculated samples, we computed the fold change for each individual variable between the two conditions. Results showed that 20 and 39 metabolites were enriched in control and AgN23 conditions, respectively (Supplementary Table S1). These metabolites were sorted based on their chemical classes, revealing a strong induction of metabolites belonging to specialised metabolism, such as indoles, flavonoids or fatty acyls (Figure 2a). A PLS-DA model was then built to identify the most significant metabolites supporting samples separation (Figure 2b). It turned out that primary metabolism markers (sucrose and glutamate) were enriched in the control root, suggesting that these are depleted from the roots in presence of the bacteria. In contrast, camalexin and indol-3-yl-methylglucosinolate (I3M) were the two most significant enriched metabolites in AgN23 treated roots. This suggests that the biosynthesis of these two metabolites is induced by the bacteria. This conclusion was further substantiated by comparing the peak areas corresponding to the two metabolites in mock and AgN23 treated plants. In the presence of AgN23, 259.4 and 2.2-fold induction were noted for camalexin and I3M, respectively (Figure 2c).

**Figure 2:**
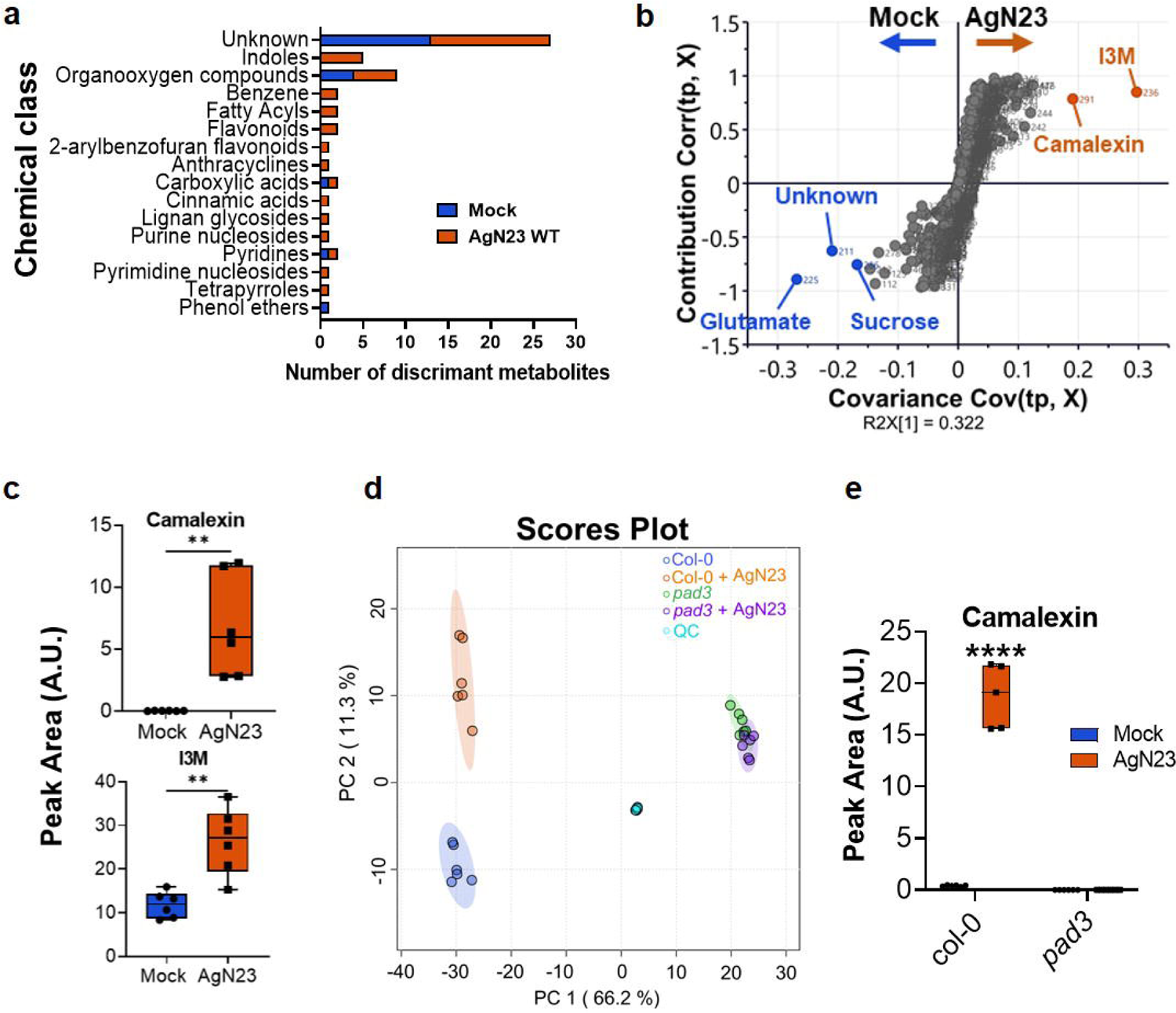
AgN23 induces camalexin biosynthesis in *Arabidopsis* roots. **a.** Discriminant metabolites overrepresented in mock or AgN23 treated *Arabidopsis* roots based on UHPLC-MS profiling data. The metabolites are displayed as chemical classes, determined with Classyfire, using the criteria of *P* value < 0.05 (T-test, control vs treatment, unadjusted *P* value) and log2 fold change (log2FC) > 0.8 or < −0.8. **b.** Corresponding S-plot of OPLS-DA score plot based on Mock vs AgN23 comparison (n = 511 variables, the OPLS-DA model was validated by a permutation test with 200 counts). The variables with VIP > 3.5 are indicated by orange and blue filled circles for AgN23 group and mock group, respectively. c. Average peak area of the 2 biomarkers significantly induced in AgN23 treated roots (VIP > 3.5). Box plots were created from data from six independent assays (n = 6). The whiskers encompass the minimum and maximum values, and the midline shows the median. Statistical differences between the treatments were analyzed using unpaired T-test and ‘**’ represents significant differences at *P* value < 0.01. I3M: indole-3-yl-methyl. **d.** PCA score plot of UHPLC-MS data (n = 534 variables) from extracts of *Arabidopsis* Col-0 or *pad3-1* 10 days after inoculation with AgN23. **e.** Average peak area of camalexin. Box plots were created from data from six independent assays (n = 6). The whiskers encompass the minimum and maximum values, and the midline shows the median.

In view of the strong and specific production of camalexin in response to AgN23, we characterised the behaviour of the *phytoalexin deficient mutant 3 (pad3-1),* mutated in a CYP450 coding gene which converts cysteine-indole-3-acetonitrile to camalexin, in response to the bacteria (Schuhegger et al 2006). Metabolomics characterisation of *pad3-1* roots indicated that the metabolome of *pad3-1* upon AgN23 inoculation was indistinguishable from that under mock conditions (Figure 2d, Supplementary Table S2). We further validated the complete lack of induction of camalexin biosynthesis in *pad3-1* (figure 2e).

Interestingly, we observed that *pad3-1* plants inoculated with AgN23 showed a phenotype similar to that of the wild type Col-0 (Figure 3a, 3b) with respect to roots and rosette growth inhibitions (Figure 3c, 3d). To check if camalexin production had any effect on AgN23 multiplication in the rhizosphere, we quantified AgN23 in the rhizosphere of the WT and the *pad3-1* mutant. Results showed that the number of genome copies of AgN23 in the rhizosphere of *pad3-1* plant was 2.96 times lower than in Col-0 (Figure 3e). Taken together with the data from *in vitro* inoculation, these results demonstrate that the induction of camalexin synthesis promotes AgN23 colonisation in the rhizosphere.

**Figure 3:**
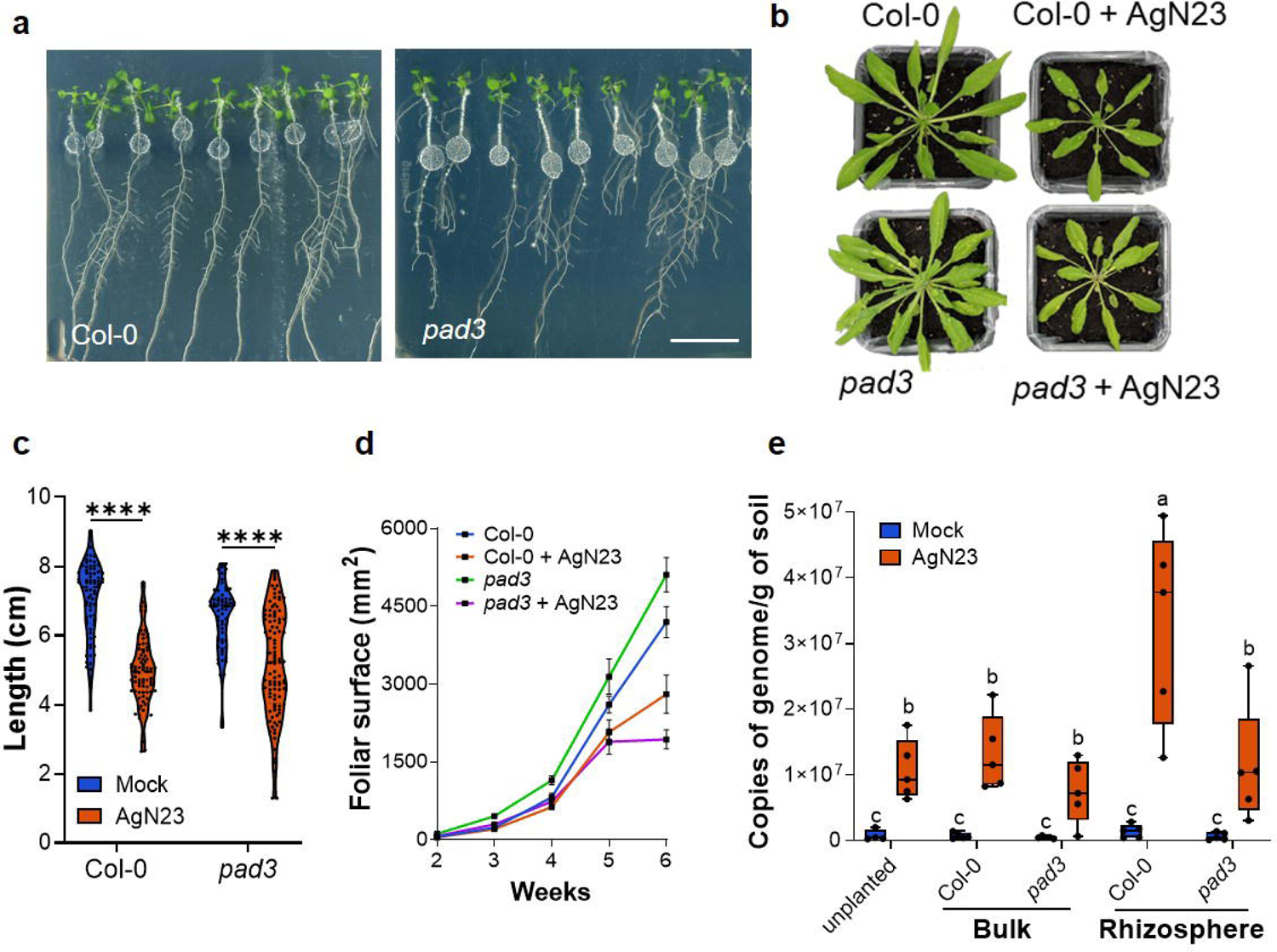
Biosynthesis of camalexin enables enrichment of AgN23 in the *Arabidopsis* rhizosphere. **a.** Observation of *A. thaliana* Col-0 and *pad3-1* colonization by AgN23 at 10 days after inoculation with spores at the root apex. Scale bar = 2 cm **b.** Rosette development of the plants Col-0 and *pad3-1* after inoculation with AgN23 spores. Typical photographs of 6-week-old col-0 or pad3 rosettes are shown. **c.** Primary root length of plants colonized or not by AgN23 at 10 days after inoculation. **d.** Leaf area measurement. Graphs show the mean ± SD calculated from at least eight biological replicates (n = 8). **e.** AgN23 genome copy number in Col-0 or *pad3-1 Arabidopsis* rhizosphere 6 weeks after soil inoculation with AgN23. Box plots were created from data from 5 plants per treatment (n = 5). The whiskers encompass the minimum and maximum values, and the midline shows the median. Letters a to c represent statistical differences between the treatments based on 2-way ANOVA followed by Tukey’s multiple comparisons test.

### AgN23 produces galbonolides, polyketides capable of inhibiting plant Inositol Phosphorylceramide synthase

To identify AgN23 specialised metabolites that could be involved in elicitation of root metabolome responses, we performed a LC-HRMS global metabolomics analysis. Briefly, apolar compounds of the CME were adsorbed on XAD16 resin beads and extracted with butanol prior to preparation for full-scan liquid chromatography high-resolution mass spectrometry (LC-HRMS). The samples were injected in ESI+ and ESI-mode and combined in a single list of variables. A total of 1022 variables were retrieved across all the samples and 812 received level 3 annotations according the Metabolomic Standard Initiative based on internally built database and the exact mass and fragmentation profile of the ions (Supplementary Table S3). This approach led to the putative identification of several specialised metabolites that have been known to be produced by *Streptomyces* ssp. (Supplementary Figure S3, Table 1).

**Table 1:**
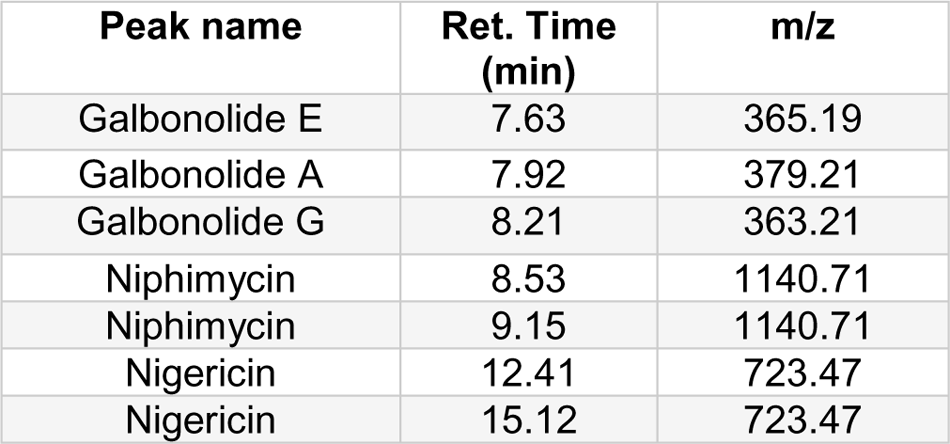
List of detected metabolites with highest intensities on chromatogram.

| Peak name | Ret. Time (min) | m/z |
| --- | --- | --- |
| Galbonolide E | 7.63 | 365.19 |
| Galbonolide A | 7.92 | 379.21 |
| Galbonolide G | 8.21 | 363.21 |
| Niphimycin | 8.53 | 1140.71 |
| Niphimycin | 9.15 | 1140.71 |
| Nigericin | 12.41 | 723.47 |
| Nigericin | 15.12 | 723.47 |

Among these specialised metabolites, we identified the antifungal compounds niphimycin, nigericin and galbonolides (also known as rustmicin). The identification of these specialised candidate metabolites is consistent with the biosynthetic gene clusters (BGCs) that we recently annotated (Gayrard et al 2023). Among the three compounds, galbonolides were originally reported for their inhibitory activities against fungal and in plants Inositol Phosphorylceramide synthase (IPCS), an enzyme involved in the metabolism of sphingolipids (Bromley et al 2003, Mandala et al 1998). The loss of function of an IPCS gene in *Arabidopsis* has been shown to be associated with programmed cell death linked to defence mechanisms (König et al 2021, Luttgeharm et al 2015, Ternes et al 2011, Wang et al 2008, Zeng et al 2022, Zienkiewicz et al 2020). Given that the inhibition of plant IPCS can trigger SA-dependent HR-like lesions, such as those observed in response to AgN23 CME, we decided to study the implication of galbonolides in *Arabidopsis* responses to AgN23 (Vergnes et al 2020). We constructed AgN23 knock out mutants in the polyketide synthase of the galbonolides BGCs by disrupting the *gbnB* gene (AS97_41300) (Figure 4a). Galbonolides detection was fully abolished in the CME of galbonolides knock-out mutants (Figure 4b). This finding confirmed the function of the predicted galbonolide gene cluster in the synthesis of all the galbonolides detected (galbonolides A, B, E and G).

**Figure 4:**
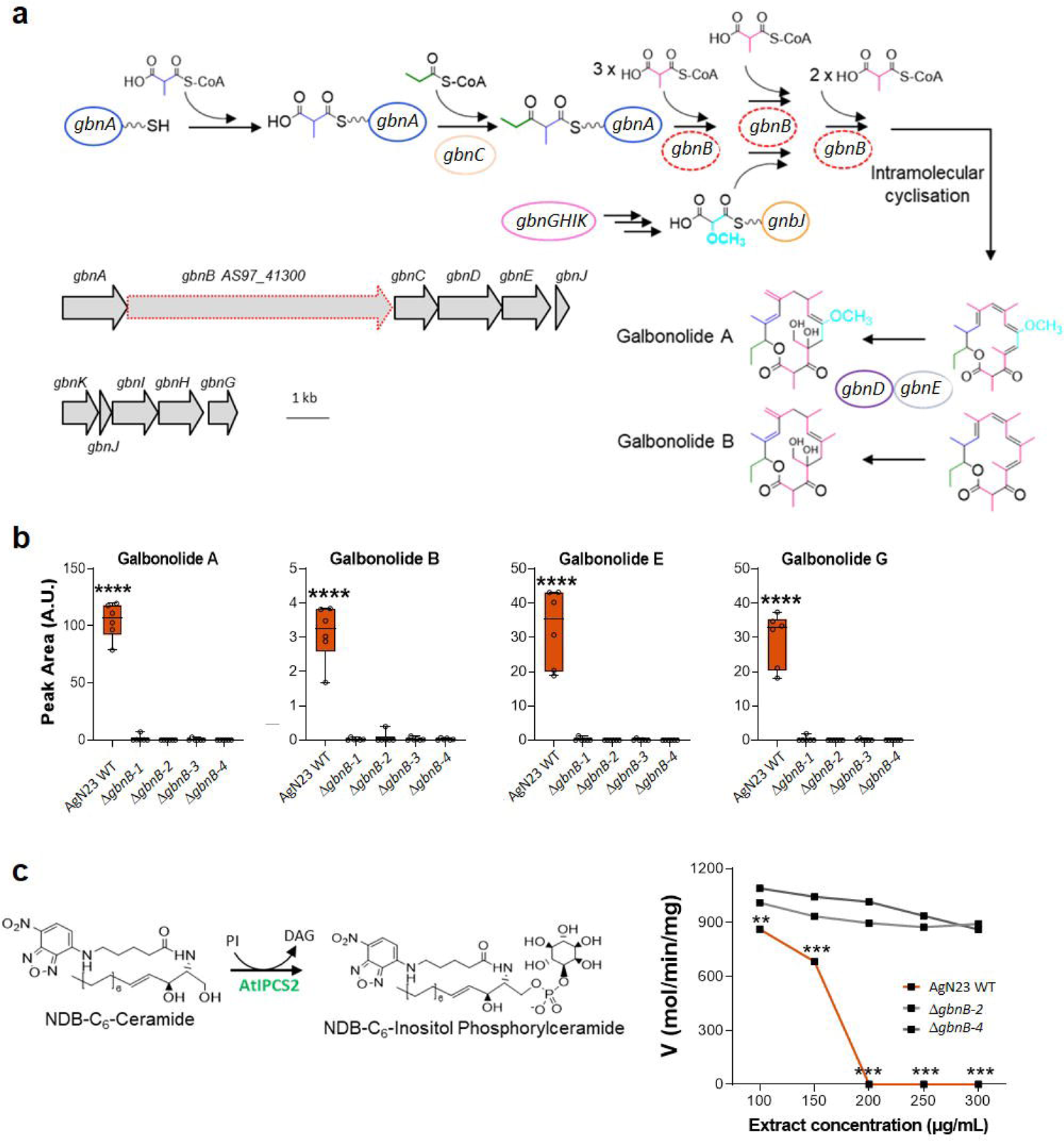
AgN23 produces galbonolides, a class of macrolides capable of inhibiting plant Inositolphosphoryl Ceramide Synthase. **a.** Biosynthesis pathway of galbonolides (*gbn*A-E) and of methoxymalonyl-CoA (*gbn*H-K) in AgN23. The targeted locus of the AgN23 galbonolide BCG for knock-out is shown by the red broken line (NCBI locus tags from assembly GCA_001598115.2). **b.** Average peak area of the different putative galbonolides structures detected in AgN23 Culture Media Extract based on HRMS and MS/MS spectra. Box plots were created from data from 6 biological replicates (n = 6). The whiskers encompass the minimum and maximum values, and the midline shows the median. Statistical differences between the AgN23 wild-type (WT) group and the AgN23 KO (Δ*gbnB*) groups were analyzed using one-way analysis of variance (ANOVA) and Tukey’s HSD test (α = 0.05) and ‘****’ represents significant differences at *P* value < 0.0001. **c.** Pathway of of NBD-C6-ceramide to NBD-C6-IPC conversion by the *Arabidopsis* Inositolphosphoryl Ceramide Synthase (AtIPCS2) and enzyme activity following treatments with butanol extracts from culture supernatant of AgN23 WT or KO mutants (Δ*gbnB-*2 and Δ*gbnB-4*). Graphs show the mean ± SD calculated from 6 independent assays (n = 6). Statistical differences between the AgN23 wild-type (WT) group and the AgN23 KO (Δ*gbnB*) groups were analyzed using multiple Mann-Whitney test (FDR = 1%) and ‘***’ and ‘**’ represent *P* value < 0.001 and *P* value < 0.01, respectively.

To investigate the effect of AgN23 and galbonolide mutants CMEs on the IPCS activity, we prepared a microsomal fraction from a *Saccharomyces cerevisiae* strain producing recombinant *Arabidopsis* IPCS2 (AT2G37940)(Pinneh et al 2019) and IPCS enzymatic activity was tracked by HPLC-Fluorescence method with the fluorescent substrate NBD-C6-ceramide (Zhong et al 1999). Data were fitted to the Michaelis-Menten equation and the apparent Km and Vmax were estimated to be 7.57µM and 0.01mol.min^-1^.mg^-1^ of protein, respectively (Pinneh et al 2019) (Supplementary Figure S4).

We then tested the enzymatic activity in presence of AgN23 and mutant CMEs in the concentration range of 100– 300µg/ml. We observed that the AgN23 CME displayed a drastic inhibition of the enzymatic activity at concentrations greater than 200µg/ml dilution (Figure 4c) whereas no such inhibition was observed in the CME of two selected AgN23 galbonolides knock-out mutants (Δ*gbnB-2* and Δ*gbnB-4*). Taken together, these data revealed that galbonolides secretion by AgN23 is the driving factor in the inhibition of *Arabidopsis* IPCS2.

Complementarily, since galbonolides were originally described as antifungal metabolites, the antifungal activity of the mutant was analysed against the filamentous fungus *Botrytis cinerea* (Harris et al 1998, Takatsu et al 1985). As expected, the loss of galbonolides in knock out mutants resulted in a reduced antifungal activity of the CME (Supplementary Figure S5, Table 2).

**Table 2:** 50% inhibitory concentration (IC_50_) of AgN23 WT and KO mutants (Δ*gbnB*) CME against *Botrytis cinerea.* Table shows mean ± SD calculated from six biological replicates (n = 6)

| Strain | IC<sub>50</sub> (μg/mL) |
| --- | --- |
| AgN23 WT | 33.28 ± 0.20 |
| $\Delta gbnB-1$ | 48.06 ± 1.68 |
| $\Delta gbnB-2$ | 47.27 ± 1.29 |
| $\Delta gbnB-3$ | 46.13 ± 1.29 |
| $\Delta gbnB-4$ | 54.42 ± 3.7 |

### Galbonolides are major contributors of the AgN23 eliciting activity and play a crucial role in rhizosphere colonisation by AgN23

In a previous work, we identified AgN23 as a *Streptomyces* strain producing strong elicitors of the hypersensitive reaction (HR) including localised necrosis and expression of defence markers such as *Pathogenesis Related 1 PR1)*, *Phytoalexin Deficient 4 (PAD4)* and *Phytoalexin Deficient 3 (PAD3*)(Vergnes et al 2020). Here, we investigated whether galbonolides may play important role in these responses to the bacterium. Agroinfiltration of *Nicotiana benthamiana* leaves with AgN23 CME induced cell death at 50 µg/ml concentration whereas no sign of necrosis could be observed at the same concentration with CMEs of the galbonolides mutants Δ*gbnB-2* and Δ*gbnB-4* (Figure 5a). It is noteworthy that similar necrotic responses were observed when CME of the wild type and mutant strains were infiltrated at 200 µg/ml or higher concentrations, suggesting that other necrotic elicitors were produced by the mutants. To investigate the effect of AgN23 CME on the necrotic responses of *Arabidopsis*, we performed ion leakage assays from infiltrated leaf discs of *Arabidopsis* with the 4 independent mutants of AgN23 and further confirmed the reduction in necrotic responses triggered by AgN23 when galbonolides biosynthesis is abolished (Figure 5b). As variation of nuclear calcium concentration is a typical signal associated with HR, we analysed the nuclear calcium concentration of *Arabidopsis* plants following treatment using a line expressing a nuclear apo-aequorin reporter gene. This reporter line was also selected based on a previous observation that a nuclear calcium signal controls the apoptotic cell death induced by d-*erythro*-sphinganine, a compound related to the sphingolipid pathway, in tobacco cells (Lachaud et al 2010). Luminescence quantification triggered by AgN23 CME in hydroponically grown *Arabidopsis* peaked at 4 minutes post treatment and this signature was abolished in the galbonolides mutants (Figure 5c). Similarly, we analysed by live imaging *Arabidopsis* seedlings inoculated at the root tip with AgN23 CME and observed that this treatment resulted in a quick activation (<15 min) of nuclear calcium signalling in the root tip which then spread to the entire root plantlets (Supplementary Movie).

**Figure 5:**
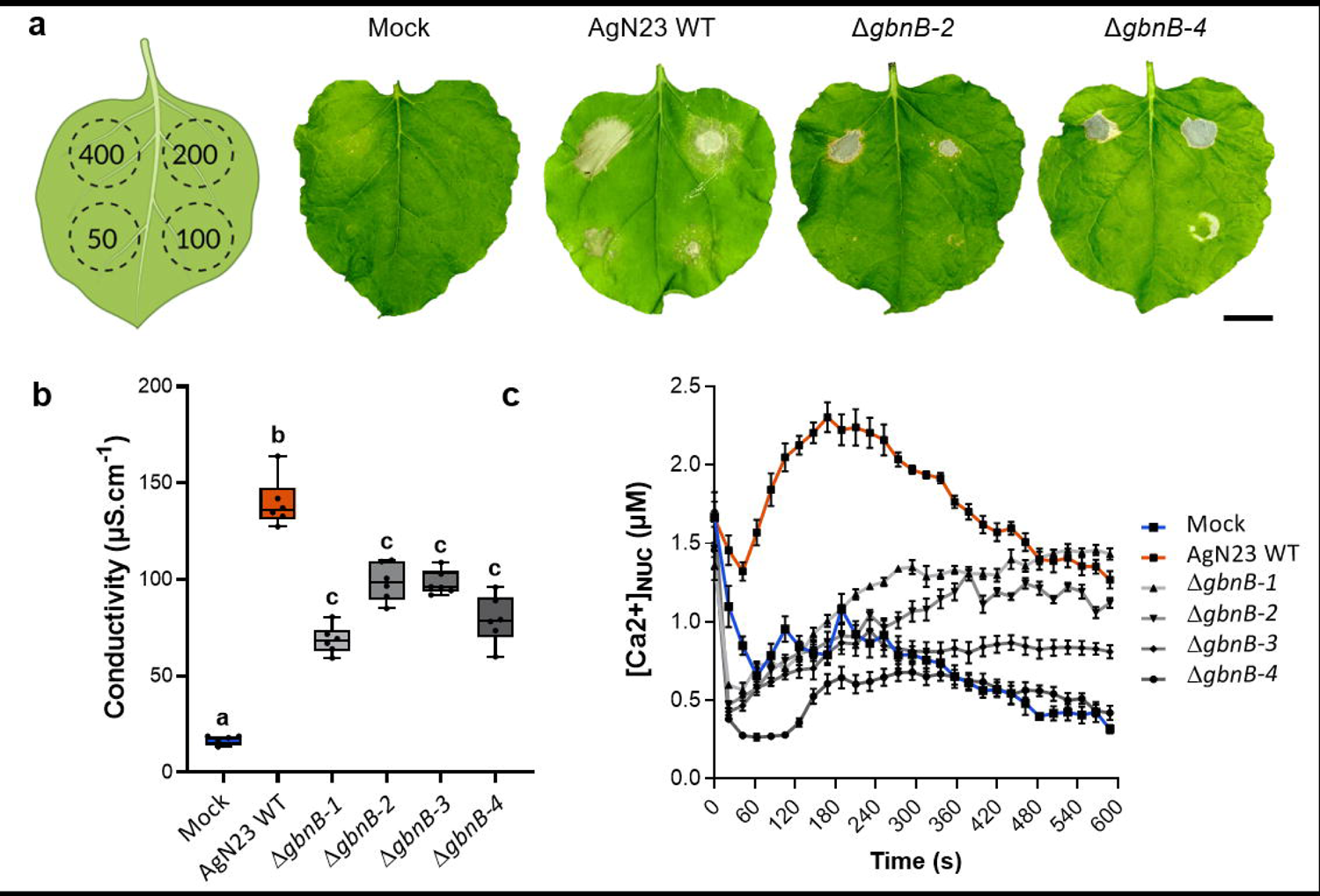
The Culture Media Extract of AgN23 triggers a galbonolides dependent hypersensitive response. **a.** Typical photographs of necrotic symptoms in *Nicotiana benthamiana* leaves 48h after infiltration with CME of AgN23 WT or KO mutants (Δ*gbnB-2* and Δ*gbnB-4*) at 50, 100, 200 and 400 µg/mL as indicated in the scheme (n = 6). Scale bar: 3 cm. **b.** Ion leakage measurements of *Arabidopsis* leaf disks infiltrated with CME of AgN23 WT or KO mutants (Δ*gbnB*) at 100 µg/mL. Box plots were created with data from 6 independent assays involving 5 to 6 leaf disks (n = 6). The letters a–c indicate statistically significant differences according to one-way analysis of variance (ANOVA) and Tukey’s HSD test (Honestly Significantly Different, α = 0.05). **c.** Kinetics of AgN23 or KO mutants CME-induced nuclear calcium influxes in *Arabidopsis seedlings expressing* nuclear-localized aequorin. CME at 100 µg/mL was added at time = 0 min. Graphs show the mean ± SD calculated from 10 independent assays involving 3 plants per treatment (n = 10).

To investigate the impact of galbonolides production on root development, *in vitro* grown seedlings were inoculated with galbonolides mutants and no root growth inhibition was observed with the two mutants (Figure 6a and 6b). Furthermore, the robust induction of expression in *PR1*, *PAD3,* and *PAD4* by AgN23 CME was compromised when using the 4 galbonolides KO mutants (Figure 6c). Thus, our data prove that galbonolides are required for the activation of immune gene expression in *Arabidopsis* seedlings in response to AgN23.

**Figure 6:**
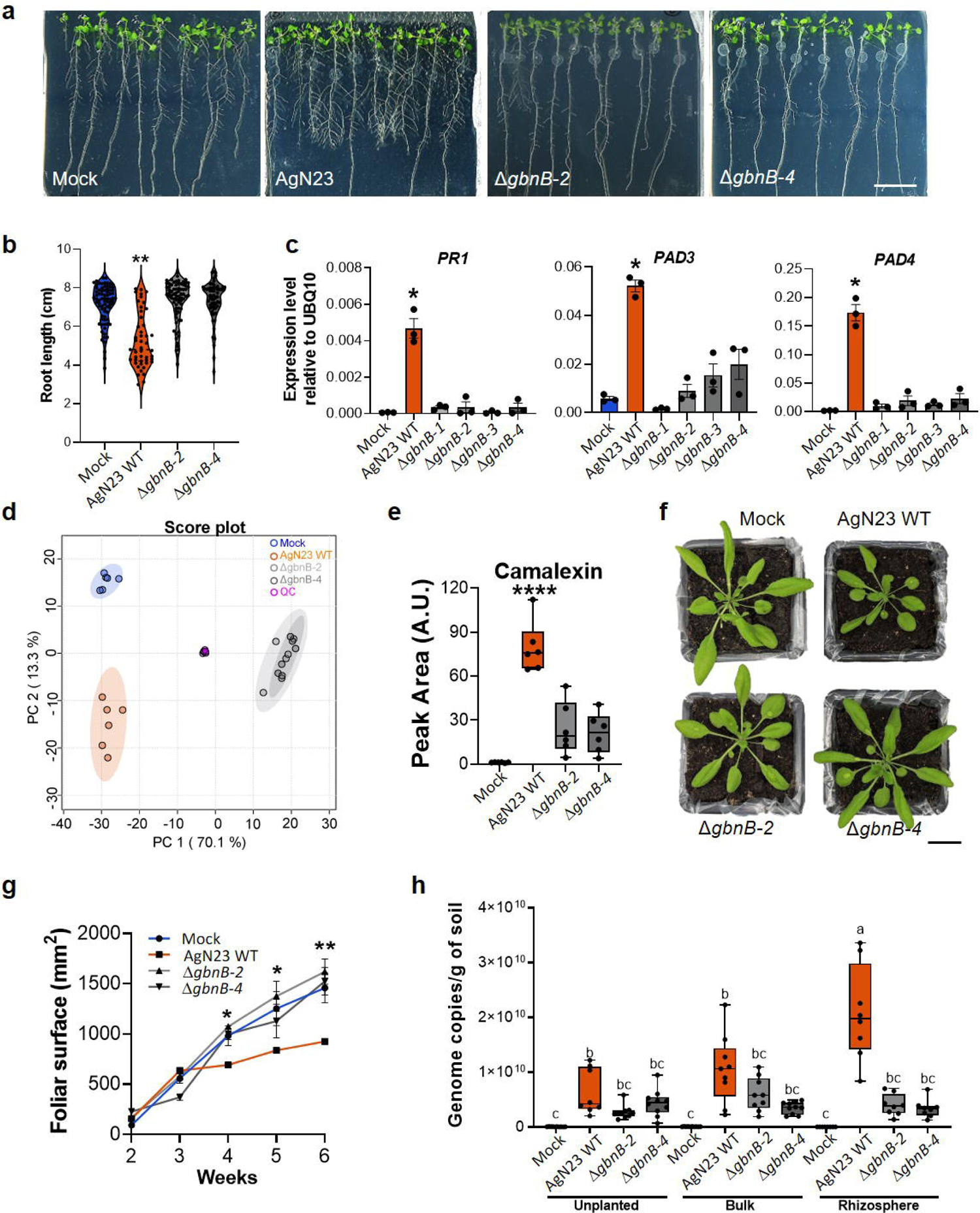
Galbonolides play a crucial role in defense gene activation, camalexin biosynthesis and AgN23 persistence in the rhizosphere. **a.** Observation of *Arabidopsis thaliana* Col-0 colonization by AgN23 WT or KO mutants at 10 days after inoculation with spores at the root apex. Scale bar: 2 cm. **b.** Primary root length of plants colonized by AgN23 WT or KO mutants (Δ*gbnB*) at 10 days after inoculation. Statistical differences between the treatments were analyzed using Mann–Whitney test and ‘****’ represents significant differences at *P* value < 0.0001. **c.** Analysis of *PR1*, *PAD3*, and *PAD4* defense gene expression in 10-day old *Arabidopsis* seedlings at 6 hours after treatment with AgN23 CME. Graphs show the mean 2^-ΔCp^ relative to *UBQ10* and SD calculated from three biological replicates (n = 3), each involving five plants. Statistical comparisons were performed with T-test (‘*’ = *P* value < 0.05). **d.** PCA score plot of UHPLC-HRMS data (n = 256 variables) from extracts of *Arabidopsis thaliana* 10 days after inoculation with AgN23 WT or KO mutants (Δ*gbnB*) **E.** Average peak area of camalexin. Box plots were created from data from six independent assays (n = 6). The whiskers encompass the minimum and maximum values, and the midline shows the median. Statistical differences between the treatments were analyzed using unpaired T-test and ‘****’ represents significant differences at *P* value < 0.0001.**f.** Typical photographs of 6-week-old Col-0 rosettes grown in potting soil inoculated with AgN23 WT or KO mutants spores. Scale bar: 1.7 cm **g.** Leaf area measurement. Graphs show the mean ± SD calculated from at least eight biological replicates (n = 8). **h.** AgN23 WT and Δ*gbnB-2* and Δ*gbnB-4)* mutants genome copy number in Col-0 rhizosphere 6 weeks after soil inoculation. Box plots were created from data from 8 plants per treatment (n = 8). The whiskers encompass the minimum and maximum values, and the midline shows the median. The letters a–c represent statistical differences between the treatments based on 2-way ANOVA followed by Tukey’s multiple comparisons test.

In view of our finding that *Arabidopsis* responds to AgN23 by strongly activating camalexin biosynthesis, we studied the effect of *in vitro* spore inoculation of Col-0 with AgN23 and the two galbonolide mutants Δ*gbnB-2* and Δ*gbnB-4*, by LC-HRMS metabolic fingerprinting (Supplementary Table S4). The PCA revealed a significant difference in the metabolome response to the two AgN23 mutants as compared to the wild type (Figure 6d). Strikingly, the induction of camalexin detection was significantly lower in roots inoculated with galbonolides mutants as compared to the wild type bacterium (Figure 6e).

Soil inoculation with galbonolides mutants did not resulted in the reduced growth of Col-0 rosette triggered by the wild-type bacteria (Figure 6f, 6g). In addition, Col-0 rhizosphere colonisation by galbonolides mutants was reduced by contrast with wild type AgN23 (Figure 6h). Put together, these data clearly point to the crucial role played by galbonolides for the induction of plant responses as well as the ability of the bacterium to colonise the rhizosphere.

## DISCUSSION

Understanding the chemical basis of the communication between plants and their associated microorganisms is essential to improve the function and composition of plant microbiota, notably in the context of developing sustainable agriculture practices. Towards this effort, *Streptomyces* species could play a major role due to their ability to efficiently colonize the rhizospheric niche and to produce a wide array of specialized metabolites with various biological activity. However, mechanisms involved in the establishment and long-term maintenance of active microbial strains in the rhizosphere are largely unknown.

To gain insight into these mechanisms we focussed on a *Streptomyces* strain, AgN23, initially isolated from the grape rhizosphere and that efficiently colonizes the rhizosphere of *A. thaliana*. Metabolic fingerprinting of the *Arabidopsis* response to AgN23 revealed that the response is mainly characterised by the production of camalexin, which is the primary *Arabidopsis* phytoalexin involved in resistance to fungal pathogens but also in the regulation of root microbiota composition and the recruitment of PGPRs (Koprivova et al 2019, Koprivova et al 2023, Nguyen et al 2022). The use of *pad3-1* camalexin deficient mutant of *Arabidopsis* demonstrated that the efficient colonization of the rhizosphere by AgN23 relies on the production of this compound. While camalexin is an antimicrobial compound, the *pad3-1* mutants did not show any signs of over colonisation by AgN23, suggesting that camalexin does not act as an inhibitor of AgN23 development but, on the reverse, favours colonisation of the rhizosphere by AgN23. Albeit camalexin is produced in response to a number of bacterial and fungal phytopathogens, this does not mean it is biologically active against these microorganisms (Glawischnig et al, 2007). It was reported that concentrations up to 500 μg.ml^-1^ are required to achieve membrane disruption in Gram negative bacteria, a range of concentration unlikely to be observed in or around *Arabidopsis* roots (Rogers et al, 1996). By contrast fungal colonisers of plant roots are sensitives to lower doses of camalexin (Rogers et al, 1996). Thus, the precise role of camalexin in supporting the development of AgN23 in the rhizosphere remains to be elucidated, but it can be hypothesized that camalexin can reduce the proliferation of susceptible fungi increasing available nutritional resources for AgN23. Recently, it has been shown that camalexin, and more generally tryptophan-derived metabolites, has been shown to be essential to prevent fungal dysbiosis in the *Arabidopsis* rhizosphere (Wolinska et al 2021).

To understand the molecular mechanisms underlying the induction of camalexin biosynthesis by AgN23, we investigated the composition of the bacteria exometabolome using untargeted metabolomic tools. This analysis identied several specialised compounds with known antimicrobial activity and for some of them, a putative function in eliciting plant defences. Among these compounds, we decided to delve into the role of galbonolides in AgN23’s biological activities, since it has been shown that galbonolides target the sphingolipid metabolism by inhibiting the Inositol Phosphorylceramide Synthase (IPCS) in both plants and fungi (Bromley et al 2003, Mandala et al 1998). Sphingolipids are signalling molecules known to play a major role in plant defence (Berkey et al 2012), notably for the activation of camalexin biosynthesis (Zienkiewicz et al 2020). Fungal toxins acting on this metabolism such as Fumonisin B1, an inhibitor of ceramide synthase produced by pathogenic *Fusarium* sp. may result in locally modifying the ceramide composition leading to induction of a hypersensitive response (Iqbal et al 2021, Zeng et al 2020). To investigate the role of galbonolides in the induction of plant defences by AgN23 we produced galbonolide mutants through the disruption of a single BGC, confirming earliest reports indicating that all galbonolide variants are produced through a single BGC (Karki et al 2010, Kim et al 2014, Liu et al 2015a, Liu et al 2015b). Using these mutants we performed a set of complementary experiments which pointed to the major requirement of galbonolides to trigger plant responses to AgN23 colonization.

Importantly, the lack of enrichment of galbonolide mutants in the *Arabidopsis* rhizosphere shows that induction of plant defence by these compounds are beneficial for the AgN23 rhizospheric lifestyle. The connections between plant immune responses and stimulation of root microorganisms has been recently exemplified (Harbort et al 2020, Singh et al 2023, Stringlis et al 2018a, Voges et al 2019). For example, the Plant Growth Promoting Rhizobacteria (PGPR) *Pseudomonas* sp. CH267 triggers the production of camalexin (Koprivova et al 2019, Koprivova et al 2023). Similarly, *Arabidopsis* root inoculation with the proteobacteria *Pseudomonas simiae* WCS417 results in the secretion of scopoletin, a coumarin that facilitates *P. simiae* root colonisation while inhibiting the growth of fungal pathogens and diverse other bacterial taxa (Stringlis et al 2018b).

However, to our knowledge, the role of a specific microbial compound in eliciting plant defense responses for the benefit of the microganism has not yet been described and this result raises an interesting question about the generality of the role of galbonolides in the rhizospheric microbiota. While IPCS and the role of sphingolipid metabolism in immune responses are ubiquitous in plants, the distribution of the galbonolide in the *Streptomyces* genus, and more generally in actinomycetes, remains to be precised. In our previous study we showed that the galbonolide biosynthetic gene cluster (BGC) is present in several species across the *S. violaceusniger* clade, to whom AgN23 belongs, which includes several rhizospheric isolates (Gayrard et al 2023). In addition, the fact that galbonolides were initially found in *S. galbus* which does not belong to the *S. violacesuniger* clade and in the distantly related actinomycete *Micromonospora spp.* suggests that the biosynthesis of galbonolides may be widespread across actinomycete representatives (Achenbach et al 1988, Karki et al 2010, Kim et al 2014, Shaffie et al 2001, Sigmund et al 1998). Further studies will aim to evaluate the impact of galbonolide production of microbiota functioning through the direct antifungal activity of these compounds and their impact on the production plant anti-fungal metabolites.

## DATA AVAILIBILITY

Datasets generated or analyzed during this study are included in this published article (and its Supplementary Table files). Raw data regarding RNAseq analysis and their complete analysis can be found on the NCBI Gene Expression Omnibus (GSE119986)(Vergnes et al 2020). The genome assembly of AgN23 is available on the NCBI “Nucleotide” repository (NZ_CP007153.1) with the chromosome sequence and the automatic annotation pipeline used to define gene models (Gayrard et al 2023). The LC-HRMS chromatograms of AgN23 CME and *Arabidopsis* are available on Zenodo repository (8421008).

## Supporting information

Supplementary movie: EM-CCD observations of calcium waves in the nuclei of an aequorin-expressing Arabidopsis seedling.

## ACKNOWLEDGEMENTS

We thank Dr. Paul Denny (Warwick University) for sharing the yeast line that allowed to purify microsomal fractions bearing *AtIPCS2,* and Dr. Pawel Bednarek for the *pad3-1* seeds. We thank Dr. Revathi Bacsa for her help in the writing of the manuscript.

## AUTHOR CONTRIBUTIONS

Clément Nicolle, Damien Gayrard, Alba Noël, Marion Hortala, Guillaume Marti, Aurélien Amiel, Sabine Grat and Aurélie Le Ru performed the research. Clément Nicolle, Jean-Luc Pernodet, Sylvie Lautru, Bernard Dumas and Thomas Rey wrote the paper.

## REFERENCE

Achenbach H, Mühlenfeld A, Fauth U, Zähner H (1988). The galbonolides. Novel, powerful antifungal macrolides from Streptomyces galbus ssp. eurythermus. Ann N Y Acad Sci 544: 128–140.

Andrić S, Meyer T, Rigolet A, Prigent-Combaret C, Höfte M, Balleux G et al (2021). Lipopeptide Interplay Mediates Molecular Interactions between Soil *Bacilli* and *Pseudomonads*. Microbiol Spectr 9: e0203821.

Andrić S, Rigolet A, Argüelles Arias A, Steels S, Hoff G, Balleux G et al (2023). Plant-associated *Bacillus* mobilizes its secondary metabolites upon perception of the siderophore pyochelin produced by a *Pseudomonas* competitor. ISME J 17: 263–275.

Avalos M, Garbeva P, Raaijmakers JM, van Wezel GP (2020). Production of ammonia as a low-cost and long-distance antibiotic strategy by *Streptomyces* species. ISME J 14: 569–583.

Berendsen RL, Vismans G, Yu K, Song Y, de Jonge R, Burgman WP et al (2018). Disease-induced assemblage of a plant-beneficial bacterial consortium. ISME J 12: 1496–1507.

Berkey R, Bendigeri D, Xiao S (2012). Sphingolipids and Plant Defense/Disease: The “Death” Connection and Beyond. Frontiers in Plant Science 3: 68.

Bromley PE, Li YO, Murphy SM, Sumner CM, Lynch DV (2003). Complex sphingolipid synthesis in plants: characterization of inositolphosphorylceramide synthase activity in bean microsomes. Arch Biochem Biophys 417: 219–226.

Bulgarelli D, Rott M, Schlaeppi K, Ver Loren van Themaat E, Ahmadinejad N, Assenza F et al (2012). Revealing structure and assembly cues for Arabidopsis root-inhabiting bacterial microbiota. Nature 488: 91–95.

Busby PE, Soman C, Wagner MR, Friesen ML, Kremer J, Bennett A et al (2017). Research priorities for harnessing plant microbiomes in sustainable agriculture. PLoS Biol 15: e2001793.

Cha JY, Han S, Hong HJ, Cho H, Kim D, Kwon Y et al (2016). Microbial and biochemical basis of a Fusarium wilt-suppressive soil. ISME J 10: 119–129.

Chaparro JM, Badri DV, Vivanco JM (2014). Rhizosphere microbiome assemblage is affected by plant development. ISME J 8: 790–803.

Conn VM, Walker AR, Franco CM (2008). Endophytic actinobacteria induce defense pathways in *Arabidopsis thaliana*. Mol Plant Microbe Interact 21: 208–218.

Cordovez V, Carrion VJ, Etalo DW, Mumm R, Zhu H, van Wezel GP et al (2015). Diversity and functions of volatile organic compounds produced by *Streptomyces* from a disease-suppressive soil. Front Microbiol 6: 1081.

Durán P, Thiergart T, Garrido-Oter R, Agler M, Kemen E, Schulze-Lefert P et al (2018). Microbial Interkingdom Interactions in Roots Promote *Arabidopsis* Survival. Cell 175: 973–983.e914.

Fitzpatrick CR, Copeland J, Wang PW, Guttman DS, Kotanen PM, Johnson MTJ (2018). Assembly and ecological function of the root microbiome across angiosperm plant species. Proc Natl Acad Sci U S A 115: E1157–E1165.

Fitzpatrick CR, Salas-González I, Conway JM, Finkel OM, Gilbert S, Russ D et al (2020). The Plant Microbiome: From Ecology to Reductionism and Beyond. Annu Rev Microbiol 74: 81–100.

Gayrard D, Nicolle C, Veyssière M, Adam K, Martinez Y, Vandecasteele C et al (2023). Genome Sequence of the *Streptomyces* Strain AgN23 Revealed Expansion and Acquisition of Gene Repertoires Potentially Involved in Biocontrol Activity and Rhizosphere Colonization. PhytoFrontiers 3(**3**): 535–547.

Getzke F, Hassani MA, Crüsemann M, Malisic M, Zhang P, Ishigaki Y et al (2023). Cofunctioning of bacterial exometabolites drives root microbiota establishment. Proceedings of the National Academy of Sciences 120: e2221508120.

Glawischnig, E. (2007). Camalexin. Phytochemistry, 68: 401–406.

Harbort CJ, Hashimoto M, Inoue H, Niu Y, Guan R, Rombolà AD et al (2020). Root-Secreted Coumarins and the Microbiota Interact to Improve Iron Nutrition in *Arabidopsis*. Cell Host & Microbe 28(**6**): 825–837.

Harris GH, Shafiee A, Cabello MA, Curotto JE, Genilloud O, Göklen KE et al (1998). Inhibition of fungal sphingolipid biosynthesis by rustmicin, galbonolide B and their new 21-hydroxy analogs. J Antibiot (Tokyo) 51: 837–844.

Iqbal N, Czékus Z, Poór P, Ördög A (2021). Plant defence mechanisms against mycotoxin Fumonisin B1. Chem Biol Interact 343: 109494.

Karki S, Kwon SY, Yoo HG, Suh JW, Park SH, Kwon HJ (2010). The methoxymalonyl-acyl carrier protein biosynthesis locus and the nearby gene with the beta-ketoacyl synthase domain are involved in the biosynthesis of galbonolides in *Streptomyces galbus*, but these loci are separate from the modular polyketide synthase gene cluster. FEMS Microbiol Lett 310: 69–75.

T. Kieser, M. J. Bibb, M. J. Buttner, K. K. Chater, D. A. Hopwood Practical streptomyces genetics (John Innes Foundation, 2000).

Kim DR, Cho G, Jeon CW, Weller DM, Thomashow LS, Paulitz TC et al (2019). A mutualistic interaction between *Streptomyces* bacteria, strawberry plants and pollinating bees. Nat Commun 10: 4802.

Kim HJ, Karki S, Kwon SY, Park SH, Nahm BH, Kim YK et al (2014). A single module type I polyketide synthase directs de novo macrolactone biogenesis during galbonolide biosynthesis in Streptomyces galbus. J Biol Chem 289: 34557–34568.

Koprivova A, Schuck S, Jacoby RP, Klinkhammer I, Welter B, Leson L et al (2019). Root-specific camalexin biosynthesis controls the plant growth-promoting effects of multiple bacterial strains. Proceedings of the National Academy of Sciences 116: 15735–15744.

Koprivova A, Schwier M, Volz V, Kopriva S (2023). Shoot-root interaction in control of camalexin exudation in *Arabidopsis*. J Exp Bot 74: 2667–2679.

Krespach MKC, Stroe MC, Flak M, Komor AJ, Nietzsche S, Sasso S et al (2021). Bacterial marginolactones trigger formation of algal gloeocapsoids, protective aggregates on the verge of multicellularity. Proceedings of the National Academy of Sciences 118: e2100892118.

Krespach MKC, Stroe MC, Netzker T, Rosin M, Zehner LM, Komor AJ et al (2023). *Streptomyces* polyketides mediate bacteria–fungi interactions across soil environments. Nature Microbiology 8: 1348–1361.

Kurth F, Mailänder S, Bönn M, Feldhahn L, Herrmann S, Große I et al (2014). *Streptomyces*-induced resistance against oak powdery mildew involves host plant responses in defense, photosynthesis, and secondary metabolism pathways. Mol Plant Microbe Interact 27: 891–900.

König S, Gömann J, Zienkiewicz A, Zienkiewicz K, Meldau D, Herrfurth C et al (2021). Sphingolipid-Induced Programmed Cell Death is a Salicylic Acid and EDS1-Dependent Phenotype in Arabidopsis fatty acid hydroxylase (fah1, fah2) and ceramide synthase (loh2) Triple Mutants. Plant Cell Physiol 63(3): 317–325.

Lachaud C, Da Silva D, Cotelle V, Thuleau P, Xiong TC, Jauneau A et al (2010). Nuclear calcium controls the apoptotic-like cell death induced by d-erythro-sphinganine in tobacco cells. Cell Calcium 47: 92–100.

Lebeis SL, Paredes SH, Lundberg DS, Breakfield N, Gehring J, McDonald M et al (2015). PLANT MICROBIOME. Salicylic acid modulates colonization of the root microbiome by specific bacterial taxa. Science 349: 860–864.

Ling, N., Wang, T., & Kuzyakov, Y. (2022). Rhizosphere bacteriome structure and functions. Nature communications, 1: 836–849.

Liu C, Zhang J, Lu C, Shen Y (2015a). Heterologous expression of galbonolide biosynthetic genes in *Streptomyces coelicolor*. Antonie van Leeuwenhoek 107: 1359–1366.

Liu C, Zhu J, Li Y, Zhang J, Lu C, Wang H et al (2015b). In Vitro Reconstitution of a PKS Pathway for the Biosynthesis of Galbonolides in Streptomyces sp. LZ35. ChemBioChem 16: 998–1007.

Lundberg DS, Lebeis SL, Paredes SH, Yourstone S, Gehring J, Malfatti S et al (2012). Defining the core *Arabidopsis thaliana* root microbiome. Nature 488: 86–90.

Luttgeharm KD, Chen M, Mehra A, Cahoon RE, Markham JE, Cahoon EB (2015). Overexpression of *Arabidopsis* Ceramide Synthases Differentially Affects Growth, Sphingolipid Metabolism, Programmed Cell Death, and Mycotoxin Resistance. Plant Physiol 169: 1108–1117.

Mandala SM, Thornton RA, Milligan J, Rosenbach M, Garcia-Calvo M, Bull HG et al (1998). Rustmicin, a potent antifungal agent, inhibits sphingolipid synthesis at inositol phosphoceramide synthase. J Biol Chem 273: 14942–14949.

Mendes R, Raaijmakers JM (2015). Cross-kingdom similarities in microbiome functions. The ISME Journal 9: 1905–1907.

Mina JG, Okada Y, Wansadhipathi-Kannangara NK, Pratt S, Shams-Eldin H, Schwarz RT et al (2010). Functional analyses of differentially expressed isoforms of the *Arabidopsis* inositol phosphorylceramide synthase. Plant Mol Biol 73: 399–407.

Nguyen NH, Trotel-Aziz P, Clément C, Jeandet P, Baillieul F, Aziz A (2022). Camalexin accumulation as a component of plant immunity during interactions with pathogens and beneficial microbes. Planta 255: 116.

Pinneh EC, Mina JG, Stark MJR, Lindell SD, Luemmen P, Knight MR et al (2019). The identification of small molecule inhibitors of the plant inositol phosphorylceramide synthase which demonstrate herbicidal activity. Sci Rep 9: 8083.

Rey T, Dumas B (2017). Plenty Is No Plague: *Streptomyces* Symbiosis with Crops. Trends Plant Sci 22: 30–37.

Rogers, E. E., Glazebrook, J., & Ausubel, F. M. (1996). Mode of action of the Arabidopsis thaliana phytoalexin camalexin and its role in Arabidopsis-pathogen interactions. Molecular Plant Microbe Interactions, 9: 748–757.

Russ D, Fitzpatrick CR, Teixeira PJPL, Dangl JL (2023). Deep discovery informs difficult deployment in plant microbiome science. Cell **186:** 4496-4513.J. Sambrook, D. Russell, Molecular Cloning: A Laboratory Manual, 3rd Revised edition (Cold Spring Harbor Laboratory Press,U.S., 2000).

Schuhegger R, Nafisi M, Mansourova M, Petersen BL, Olsen CE, Svatoš A et al (2006). CYP71B15 (PAD3) Catalyzes the Final Step in Camalexin Biosynthesis. Plant Physiology 141: 1248–1254.

Shafiee A, Harris G, Motamedi H, Rosenbach M, Chen T, Zink D et al (2001). Microbial hydroxylation of rustmicin (galbonolide A) and galbonolide B, two antifungal products produced by *Micromonospora* sp. Journal of Molecular Catalysis B: Enzymatic 11: 237–242.

Sigmund JM, Hirsch CF (1998). Fermentation studies of rustmicin production by a *Micromonospora* sp. J Antibiot (Tokyo) 51: 829–836.

Singh G, Agrawal H, Bednarek P (2023). Specialized metabolites as versatile tools in shaping plant– microbe associations. Molecular Plant 16: 122–144.

Stringlis IA, Proietti S, Hickman R, Van Verk MC, Zamioudis C, Pieterse CMJ (2018a). Root transcriptional dynamics induced by beneficial rhizobacteria and microbial immune elicitors reveal signatures of adaptation to mutualists. Plant J 93: 166–180.

Stringlis IA, Yu K, Feussner K, de Jonge R, Van Bentum S, Van Verk MC et al (2018b). MYB72-dependent coumarin exudation shapes root microbiome assembly to promote plant health. Proceedings of the National Academy of Sciences 115: E5213–E5222.

Ternes P, Feussner K, Werner S, Lerche J, Iven T, Heilmann I et al (2011). Disruption of the ceramide synthase LOH1 causes spontaneous cell death in *Arabidopsis thaliana*. New Phytologist 192: 841–854.

Thiergart T, Durán P, Ellis T, Vannier N, Garrido-Oter R, Kemen E et al (2020). Root microbiota assembly and adaptive differentiation among European *Arabidopsis* populations. Nat Ecol Evol 4: 122–131.

Vergnes MS, Gayrard MD, Veyssiere MM, Toulotte MJ, Martinez MY, Dumont DV et al (2020). Phyllosphere colonisation by a soil *Streptomyces* sp. promotes plant defense responses against fungal infection. Molecular Plant-Microbe Interactions, 33(**2**): 223–234.

Viaene T, Langendries S, Beirinckx S, Maes M, Goormachtig S (2016). *Streptomyces* as a plant’s best friend? FEMS Microbiol Ecol 92.

Voges MJEEE, Bai Y, Schulze-Lefert P, Sattely ES (2019). Plant-derived coumarins shape the composition of an *Arabidopsis* synthetic root microbiome. Proceedings of the National Academy of Sciences 116: 12558–12565.

Wang W, Yang X, Tangchaiburana S, Ndeh R, Markham JE, Tsegaye Y et al (2008). An inositolphosphorylceramide synthase is involved in regulation of plant programmed cell death associated with defense in *Arabidopsis*. Plant Cell 20: 3163–3179.

Wolinska KW, Vannier N, Thiergart T, Pickel B, Gremmen S, Piasecka A et al (2021). Tryptophan metabolism and bacterial commensals prevent fungal dysbiosis in *Arabidopsis* roots. Proceedings of the National Academy of Sciences 118: e2111521118.

Yang Z, Qiao Y, Konakalla NC, Strøbech E, Harris P, Peschel G et al (2023). *Streptomyces* alleviate abiotic stress in plant by producing pteridic acids. Nature Communications 14: 7398.

Zeng H-Y, Bao H-N, Chen Y-L, Chen D-K, Zhang K, Liu S-K et al (2022). The Two Classes of Ceramide Synthases Play Different Roles in Plant Immunity and Cell Death. Frontiers in Plant Science 13: 824585.

Zeng HY, Li CY, Yao N (2020). Fumonisin B1: A Tool for Exploring the Multiple Functions of Sphingolipids in Plants. Front Plant Sci 11: 600458.

Zhong W, Murphy DJ, Georgopapadakou NH (1999). Inhibition of yeast inositol phosphorylceramide synthase by aureobasidin A measured by a fluorometric assay. FEBS Letters: 4.

Zienkiewicz A, Gömann J, König S, Herrfurth C, Liu YT, Meldau D et al (2020). Disruption of *Arabidopsis* neutral ceramidases 1 and 2 results in specific sphingolipid imbalances triggering different phytohormone-dependent plant cell death programmes. New Phytol 226: 170–188.

