## Supplementary Figure S1-5 for "Root-associated *Streptomyces* produce galbonolides to modulate plant immunity and promote rhizosphere colonisation"

**AFFILIATIONS**

**SHORT TITLE**

Galbonolides from rhizosphere *Streptomyces*

**KEY WORDS**

Rhizosphere, Galbonolides, *Streptomyces*, *Arabidopsis*, Camalexin

**a**

| TAIR10 Id | Name | Annotation | AgN23 1 hpi | AgN 23 6 hpi |
|---|---|---|---|---|
| At5g60890 | ATMYB34 | MYB transcription Factor | -1.689 | -1.77 |
| At1g18570 | MYB51 | MYB transcription Factor | 4.54 | 5.32 |
| At1g74080 | ATMYB122 | MYB transcription Factor | 12.378 | 14.321 |
| At4g39950 | CYP79B2 | Cytochrome P450 | -1.093 | 5.684 |
| At4g31500 | CYP83B1 | Cytochrome P450 | -1.075 | 2.754 |
| At2g30750 | CYP71A12 | Cytochrome P450 | 18.592 | 55.686 |
| At2g30770 | CYP71A13 | Cytochrome P450 | -1.294 | 153.13 |
| At3g26830 | PAD3 | Cytochrome P450 | 28.805 | 29.805 |

**b**

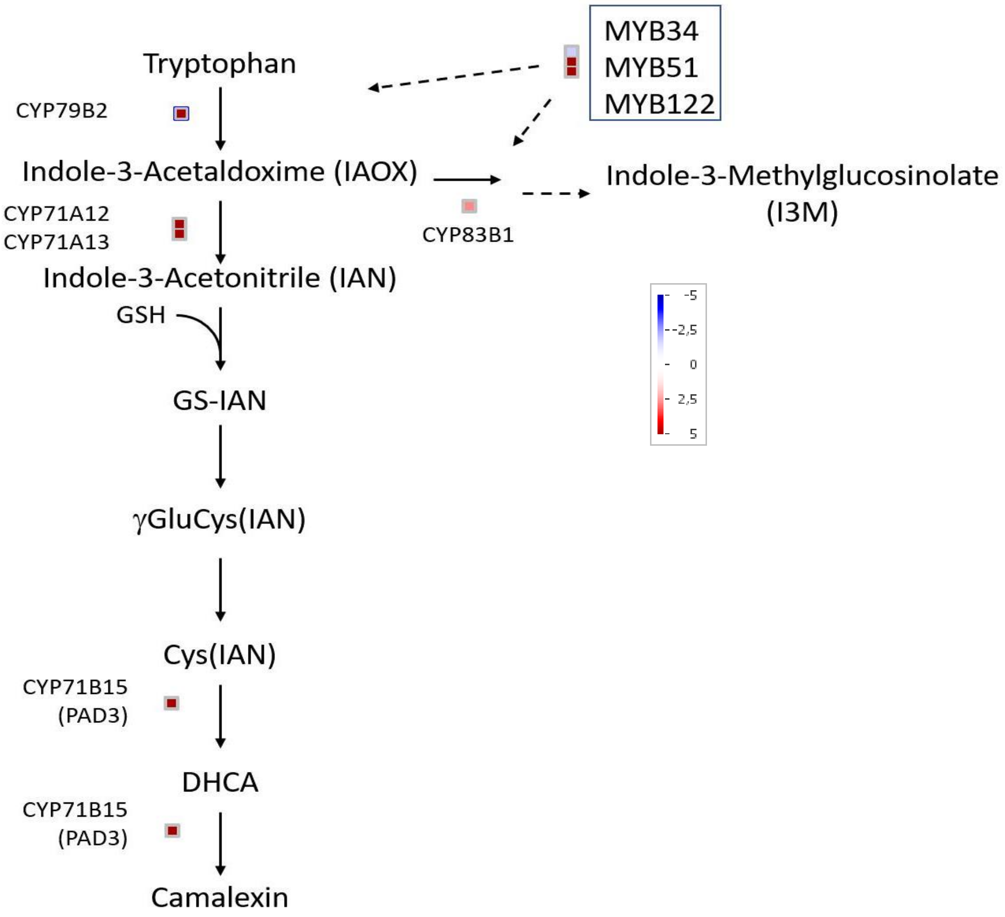


**Supplementary Figure S1:** Expression of genes involved in the biosynthesis of indolic compounds following treatment of *Arabidopsis* seedlings with AgN23 CME. Transcriptomic data from Vergnes et al., 2020 were mined to extract expression of genes falling in the category of indolic biosynthesis. **a.** Fold induction or repression expressed in Log2 of genes involved in gene regulation (MYB transcription factors) or camalexin biosynthesis at 1 hour post inoculation (hpi) and 6 hpi. **b.** Mapman display of gene regulation at 6 hpi from the biosynthesis pathway of camalexin and I3M (adapted from Ferigmann et al., MPMI Vol. 34, No. 5, 2021, pp. 560–570).

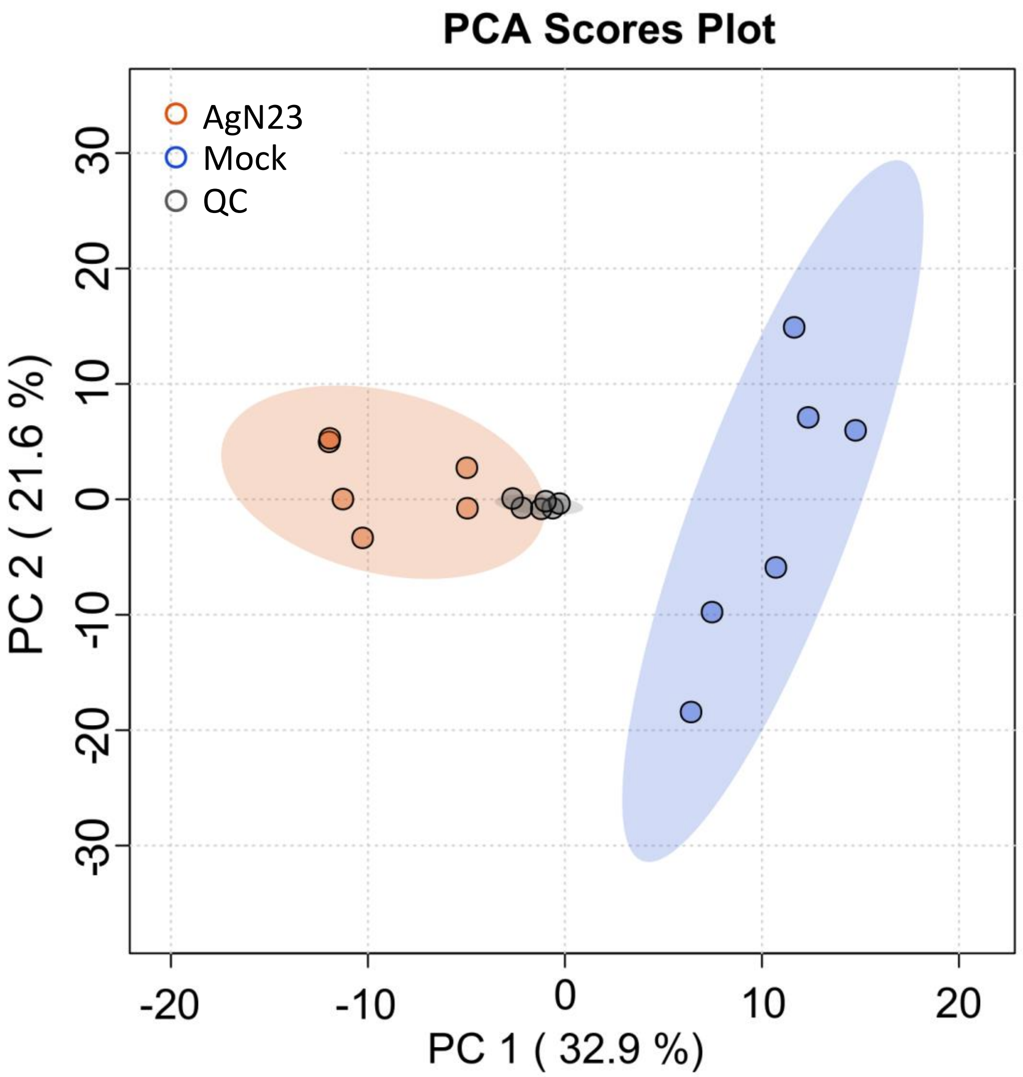


**Supplementary Figure S2: PCA score plot of UHPLC-HRMS data (n = 511 variables) from extracts of *Arabidopsis thaliana* 10 days after inoculation or not with AgN23 spores.** QC: quality control

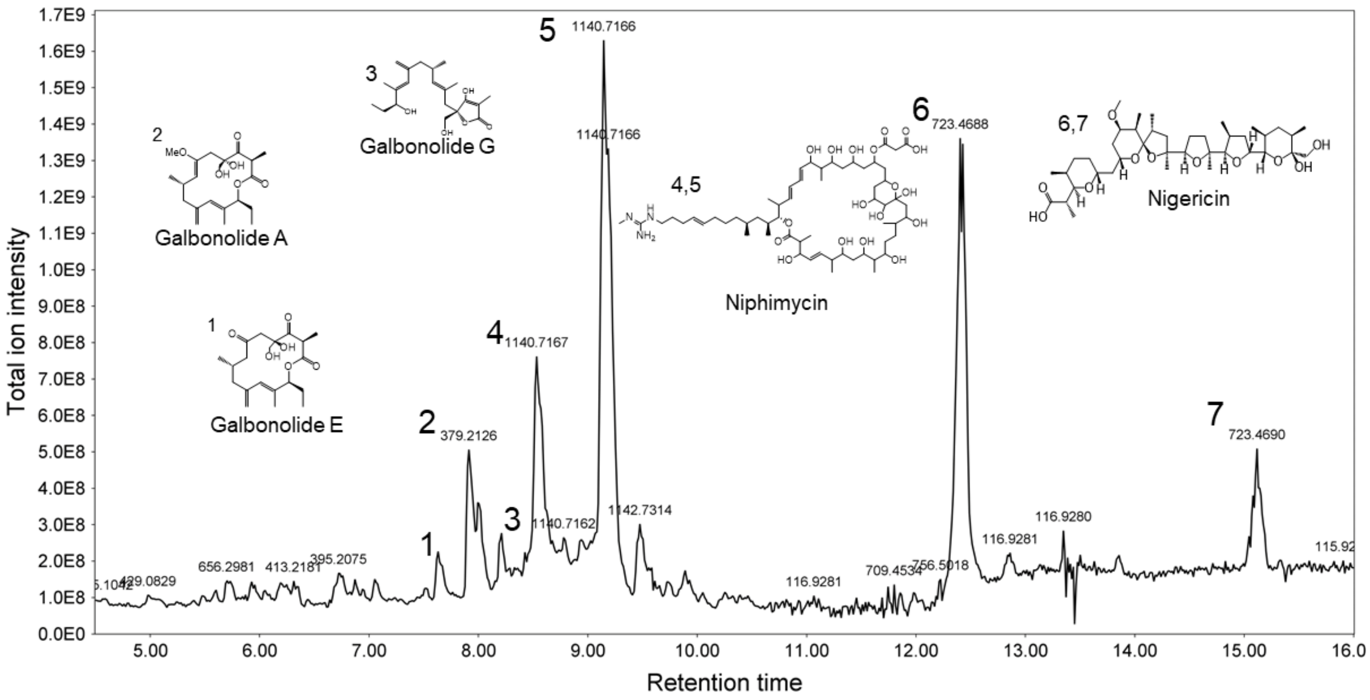


**Supplementary Figure S3:** UHPLC-HRMS chromatogram of AgN23 CME expressed in Total Ion Intensity. Peaks with the highest intensities were annotated with putative structures based on HRMS and MS/MS spectra.

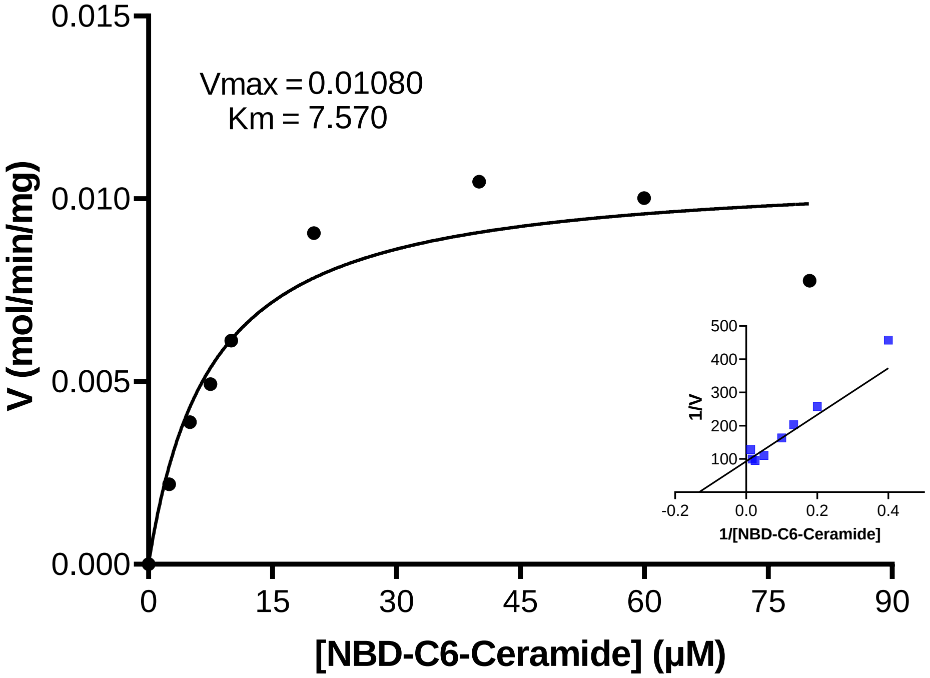


**Supplementary Figure S4: Michaelis-Menten and Lineweaver Burk Vmax, and Km estimations from enzyme assays of the Inositol Phosphoceramide synthase from Arabidopsis thaliana (AtIPCS2).** 0.1 mg/mL of total microsomal membranes were used to study the conversion of and the NBD-C6-Ceramide to NBD-C6-IPC. The fluorescence values of the assays were converted to concentrations based on the line of best fit from the standard curve of NBD-C6-Ceramide (3–500 μM).

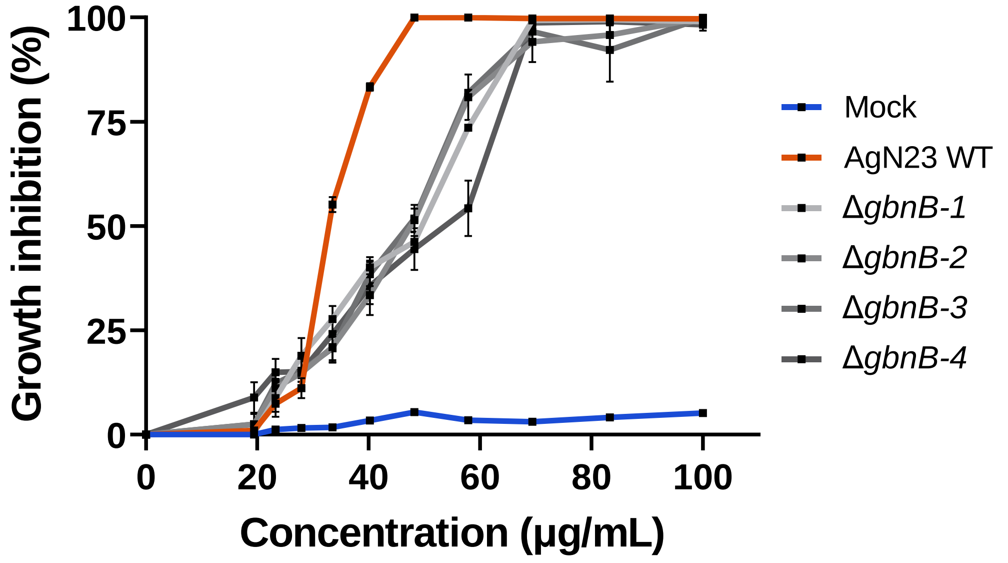


**Supplementary Figure S5: Growth inhibition of Botrytis cinerea following treatment with CME of AgN23 WT and Δ*gbnB* mutants.** Graphs show the mean ± SD calculated from six biological replicates (n = 6).
