## Supplementary methods for "Root-associated *Streptomyces* produce galbonolides to modulate plant immunity and promote rhizosphere colonisation"

**AFFILIATIONS**

**SHORT TITLE**

Galbonolides from rhizosphere *Streptomyces*

**KEY WORDS**

Rhizosphere, Galbonolides, *Streptomyces*, *Arabidopsis*, Camalexin

### SUPPLEMENTARY METHODS

#### *Arabidopsis* cultivation *in vitro*

For *in vitro* cultivation, seeds of *Arabidopsis* were surface sterilized with a mixture containing 25% NaClO and absolute ethanol in the volume ratio 1:5 for 10 minutes and then rinsed two times with 95% ethanol. Ethanol was discarded and the seeds were allowed to dry completely. Seeds were then sown onto Murashige and Skoog (MS, Sigma) at 4.4 g/L supplemented with agarose (Sigma) 8 g/L and adjusted to pH 5.7. Surface sterilized seeds were allowed to sprout in a growth phytotronic chamber (16 hours photoperiod, 23°C) on vertically placed plates filled with MS medium amended with 1 % sucrose. 2 days after germination, seedlings were transferred to 12 cm2 square Petri dishes containing 1x sucrose-free MS medium at the rate of 10 seedlings per dish. Four-day old seedlings were then inoculated at the root tip with 10 µL of spore inoculum at $10^5$ CFU/mL and placed back vertically in the growth phytotronic chamber. 10 µL of sterilized ultrapure water was used for mock treatments. Root phenotype was observed 10 days after inoculation and photographs were taken with an Expression 11000 XL scanner (Epson) at 300 dots/inch. Primary root lengths were measured with Image J software (v. 1.51k).

#### Detection of cell death related loss of electrolytes in *Arabidopsis*

The loss of electrolytes from dying cells due to hypersensitive response related programmed cell death (HR-PCD) was monitored according to a vacuum-based infiltration method of leaf material as described previously (Johansson et al 2015). Briefly, *Arabidopsis* leaf discs, prepared with a cork borer (5mm diameter), were covered in 10mL of bacterial culture supernatant extract at 100 µg dry extract/mL and placed in a SpeedVac vacuum concentrator (Thermo ScientificTM). After infiltration, the discs were poured into a tea strainer, rinsed with ultrapure water, and transferred to 6-well cell cultivation plates filled with 10 mL ultrapure water (4–5 discs in each well in 6 replicates per treatment). Conductivity was measured by transferring 8mL of the bathing solution to a 15 mL centrifuge tube in which an InLab 731-ISM conductivity cell (Mettler Toledo) was placed. The solution was returned to the plate well after measurement. Plates were kept in a growth phytotronic chamber (16 hours photoperiod, 23°C) between measurements.

**Detection of *Arabidopsis* $Ca^{2+}$ signals**

To perform nuclear calcium influx studies, we used an *Arabidopsis* transgenic line expressing the chimeric construct containing the 35S promoter controlling the nucleoplasmin coding region in frame with the coding region of apo-Aequorin as described previously (Pauly et al 2001). The vector was transformed into the *Agrobacterium tumefaciens* LBA4404 strain and Col-0 transgenic plants carrying the 35S::apo-aequorin-nuc were generated using the floral dip method and selected in 50 µg/mL hygromycin B (Clough and Bent 1998). Surface sterilized *Arabidopsis thaliana* 35s::apo-aequorin-nuc were cultivated on vertically placed plates filled with ½ MS medium amended with 1% sucrose and with a sterile 120 × 120 mm Nytex fabric (mesh opening 37 µm; Sefar) layered on top. Three 10-day old seedlings were sampled and placed in 11 x 55 mm 3 mL polystyrene tubes and considered as one replicate. For gene expression assays, Col-0 seeds were surface sterilized and transferred into the wells of 24-well plates containing 300 µL ½ MS + 1% sucrose medium. Seeds were allowed to sprout in a growth chamber (16 hours photoperiod, 23°C) mounted on a rotary shaker at 90 rpm for 2 days. 5 seedlings per well were kept and the plates were transferred on a static tray. ½ MS + 1% sucrose medium was replaced with 300 µL of fresh medium 7 days after germination. The seedlings were then treated with 300µL of CME at 200 µg dry extract/mL added to each well 10 days after germination. Sampling was performed at 1, 6, and 24 hours after treatment, the seedlings being immediately thrown into liquid nitrogen and stored at -80 °C until RNA extraction.

$Ca^{2+}$ signals in the nuclei of *Arabidopsis thaliana* cells were quantitatively measured as described previously (Mithöfer et al 2009). Holoaequorin was reconstituted by adding 400 µL of 2.5 µM coelenterazine (Interchim) solution to the tube followed by overnight incubation in darkness. At time 0, the coelenterazine solution was removed and the tube was placed in a Sirius single tube luminometer (Berthold Technologies). A volume of 400 µL of AgN23 WT or AgN23 KO mutants supernatant culture extracts at 100 µg dry extract/mL were then gently added on the seedlings and photon counting was immediately started by closing the luminometer chamber. After the $Ca^{2+}$ response reached the initial baseline level, 300 µL of lysis solution (100mM CaCl2 in 10% ethanol supplemented with 2% Nonidet P40 (v/v)) were automatically injected in the polypropylene tube to discharge the free aequorin owing to the calcium-enriched medium. The emitted light was calibrated in terms of $Ca^{2+}$ concentrations using a previously described method (Allen et al 1977).

$$[Ca^{2+}] = \frac{(\frac{L_0}{L_{max}})^{1/3} + \left[KTR \times \left(\frac{L_0}{L_{max}}\right)^{\frac{1}{3}}\right] - 1}{KR \times \left(\frac{L_0}{L_{max}}\right)^{\frac{1}{3}}}$$

L0 is the luminescence intensity/s and Lmax is the total amount of luminescence present in the entire sample over the course of the experiment. $[Ca^{+}]$ is the calculated $Ca^{2+}$ concentration, KR is the dissociation constant for the first $Ca^{2+}$ ion to bind, and KTR is the binding constant of the second $Ca^{2+}$ ion to bind to aequorin. The luminescence data were determined using the KR and KTR values of 7 × 106 M−1 and 118, respectively. For time course imaging, 5-day old *Arabidopsis thaliana* 35s::Aequorin-nuc seedlings were placed in 60mm petri dishes filled with ½ MS medium. Holoaequorin was reconstituted by adding 500 µL of 2.5 µM coelenterazine (Interchim) solution to the dish followed by overnight incubation in darkness. At time 0, the coelenterazine solution was removed and plants were treated by adding 10 µL of supernatant culture extracts at 100 µg dry extract/mL at the root apex. Images were acquired with a homemade system based on a EM-CCD camera (C9100-13, Hamamatsu) driven by HC-image software (Hamamatsu) and a macro objective. Both system and sample were placed in a dark room during the experiment. The camera was set to EM-CCD mode with the gain set at 5 and 1200, and the exposure time was 5 seconds. Images were analysed with the ImageJ software package (http://rsb.info.nih.gov/ij/; v. 1.51k).

**Analysis of *Arabidopsis* AtIPCS2 enzymatic activity**

The Initial velocities of the AtIPCS2 reaction were measured at various NBD-C6-phytoceramide (2.5 – 80 µM, final concentration, Cayman) with a fixed soybean PI concentration (2 mM, final concentration,

Sigma) and a total microsomal proteins concentration of 0.1 mg/mL. Enzymatic reactions (50 µL final volume) were done in 2 mL microtubes and incubated at 30 °C for 30 minutes. Following incubation, the reactions were quenched with 150 µL methanol. After centrifugation at 10,000 g for 3 minutes, the organic phase was removed, dried under nitrogen flow, and re-suspended in 100 µL methanol. 15 µL quantities were injected using an automatic sample injector into the HPLC instrument. The fluorescent substrate and product (Vex = 465 nm, Vem = 530 nm) were separated on a XBridge C18 reversed-phase column (Waters) with the following gradient at 1mL/min: 50% CH3CN – 50% H2O – 0.1% CH3COOH to 90% CH3CN – 10% H2O – 0.1% CH3COOH. Two HPLC peaks were observed for the substrate and reaction product at 13.8 and 4.1 minutes, respectively. 15 µL quantities of NBD-C6-phytoceramide at known concentrations were injected to the calculate AtIPCS2 velocity. As the substrate fluorescence is equal to the product fluorescence, the enzyme velocity was determined as the rate of product formation over time.

**Quantification of AgN23 genome copies from rhizosphere samples**

AgN23 specific primers were designed to quantify the AgN23 genome copy number by qPCR analysis. Briefly, 10 kilobases portions of the AgN23 genome gapless assembly (GCF_001598115.1) was blasted with NCBI Primer-BLAST with default settings except for PCR product size (70-200 bp), database (nr) and organism (Streptomyces taxid:1883). Primers AgN23-F (5'-CATGGGTTTCTGTGGCCTCT-3') and AgN23-R (5'-AGATGGTTCACGCCCACATT-3') were found and validated experimentally to target a unique intergenic region (2108022-2108187) and resulted in a 166-bp long PCR product with Tm = 59.96 °C (Data S5). qPCR was performed in triplicate using a CFX Opus Real-Time PCR System (Biorad). Each qPCR reaction was performed in 10 µL with the LightCycler® 480 SYBR Green I Master mix (Roche), 300 nM of each primer, and the template DNA. 4 ng of total DNA extracted from soil samples was used. The amplification conditions were as follows: preheating at 95 °C for 5 minutes followed by 40 cycles of 95 °C for 15 seconds and 60 °C for 60 seconds. Melting curves were checked to confirm purity of the amplified product. AgN23 pure genomic DNA was isolated as described previously (Gayrard et al 2023), quantified, and serial dilutions were performed to determine AgN23 genome concentration in soil samples. The number of gene copies per g of soil (N) was calculated as:

$$m_{AgN23} = \frac{S_{AgN23} \times M_{bp}}{Na}$$

$$N = \frac{D_{well} \times C_{sample} \times 50{,}000}{m_{AgN23}}$$

Where mAgN23 is the mass of one genome copy of AgN23 in ng (1.08.10-5 ng), SAgN23 is AgN23 genome length in base (SAgN23 = 10.9 Mb), Mbp is the molecular mass of one base pair (600 g/mol), Na the Avogadro constant (Na = 6.022.1023 $mol^{-1}$), Dwell is the DNA mass calculated in a well of qPCR reaction in ng, and Csample is the total DNA concentration of 100 mg of soil sample in ng/µL.

**Production of AgN23 spores inoculum**

To produce spore inoculum, SFM plates were incubated for two weeks at 28°C in darkness before filling them with 10 mL of sterile ultrapure water. The mycelium was thoroughly scraped with a sterile spreader and the resulting solution was vortexed, then filtered in a 20 mL syringe filled with sterile cotton wool. The resulting suspension was centrifuged at 4,200 rpm for 10 minutes. The supernatant was discarded, the pellet re-suspended in sterile ultrapure water, and the resulting suspension was adjusted to $10^5$ CFU/mL.

**Antifungal activity**

For the determination of AgN23 CME $IC_{50}$ against fungi, the *Botrytis cinerea* BD90 strain was cultivated onto solid Potato Dextrose Agar (39 g/L, Sigma). Fungal spores were harvested 8 days after cultivation by pouring 5 mL of sterile ultrapure water into the plate and gently scrapping the surface with a sterile spreader. Spores were counted on a Fuchs-Rosenthal cell and the spore suspension was adjusted to 5555 spores/mL in Potato Dextrose Broth (5.3 g/L, Sigma). 90 µL of spore suspension were distributed in 96-well plates. A volume of 10 µL of culture supernatant extracts from AgN23 WT or Δ*gbnB* mutants at different concentrations (20–100 µg dry extracts/mL) were added with 3 replicates per concentration. *B. cinerea* growth was monitored by $OD_{600}$ readings with an ELx808TM Incubating Absorbance Microplate Reader (Bio-Tek) 0, 3, and 6 days after treatment. Growth inhibition (I) was calculated as:

$$I = \frac{OD_T - OD_{T0}}{OD_{TControl} - OD_{T0Control}}$$

Where $OD_T$ is $OD_{600}$ of treatment condition at time = T, $OD_{T0}$ is $OD_{600}$ of treatment condition at time = T0, $OD_{TControl}$ is $OD_{600}$ of control condition at time = T, $OD_{T0}$ is $OD_{600}$ of control condition at time = T0.

**Biochemical preparation of *Arabidopsis* and AgN23 samples**

*Arabidopsis* flash frozen samples were ground 2 times *via* bead beating with a mixer mill (Retsch) at 30 Hz for 30 seconds. 100 mg of plant powder were placed into lysing matrix D 2 mL tubes (MP Biomedicals) containing 1 mL of extraction buffer (CH3OH: C3H8O: H2O: $CH_2O_2$ = 40: 40: 19.5: 0.5) ground twice for 20 seconds with a FastPrep-24TM homogenizer (MP Biomedicals) at 6.5 m/s. Samples were re-frozen in liquid nitrogen between each cycle. The supernatants were isolated by centrifugation at 10,000 g for 20 minutes and concentrated in a Speedvac vacuum concentrator (Thermo Scientific). Samples were then resuspended in 1 mL of injection buffer (50 % CH3OH 50% $H_2O$) and filtrated through 750 µL nonsterile micro-centrifugal filters (PTFE; 0.2 µm; Thermo Scientific) and introduced in HPLC certified vials. An aliquot of each sample from the same extraction series was pooled together for quality control (QC).

To retrieve AgN23 Culture Media Extract (CME) from the culture, Amberlite XAD-16 (Sigma) was added to each culture flask (60 g/L) and mixed overnight at 250 rpm and 28 °C in a shaking incubator. Subsequently, XAD-16 was separated through vacuum filtration and the liquid was discarded. The separated XAD-16 was then placed in 50 mL butanol for 4 hours and flasks were regularly hand-shaken. One more vacuum filtration was then carried out, the XAD-16 was discarded, and the resulting butanol extracts were then concentrated by rotary evaporation with a Rotavapor® R-300 (Buchi). The concentrated extracts were placed in calibrated 5mL test tubes, completely dried under nitrogen flow, weighed, and adjusted to 2 mg/mL in 50% $CH_3CN$ 50% H2O mixture. Finally, 700 µL quantities of the diluted extract were filtrated through 750 µL nonsterile micro-centrifugal filters (PTFE; 0.2 µm; Thermo Scientific) and introduced in HPLC certified vials. An aliquot of each sample from the same extraction series was pooled together for quality control (QC). Parameters UHPLC-HRMS profiling, feature annotations and statistical analysis are described in Supplementary methods.

**Ultra High Performance Liquid Chromatography High Resolution Mass Spectrometry analysis**

Ultra-High-Performance Liquid Chromatography-High Resolution MS (UHPLC-HRMS) analyses were performed on a Q Exactive Plus quadrupole mass spectrometer equipped with a heated electrospray probe (HESI II) coupled to a U-HPLC Ultimate 3000 RSLC system (Thermo Fisher Scientific, Hemel Hempstead, UK). Samples were separated on a Luna Omega Polar C18 column (150×2.1 mm i.d., 1.6 µm, Phenomenex, Sartrouville, France) equipped with a guard column. The mobile phase A (MPA) was H2O with 0.05% formic acid (FA) and the mobile phase B (MPB) was $CH_3CN$ with 0.05% FA. The solvent gradient started with 2% B for 30 seconds, reaching 70% B at 10.5 minutes and 98% at 10.6 minutes, holding 98% for 2 minutes, and coming back to the initial condition of 2% B in 0.1 minutes, for a total run time of 14 minutes. The flow rate was $0.3 mL.min^{-1}$, the column temperature was set to 40 °C, the autosampler temperature was set to 10 °C, and the injection volume was fixed at 2 µL. Mass detection was performed in positive ionization (PI) and negative ionization (NI) modes at 30 000 resolving power [full width at half maximum (FWHM) at 400 m/z] for MS1 and 17 500 for MS2 with an automatic gain control (AGC) target of $10^{-5}$. Ionization spray voltages were set to 3.5 kV (for PI) and 2.5 kV (for NI) and the capillary temperature was set to 256°C for both modes. The mass scanning range was m/z 100-1500 Da. Each full MS scan was followed by data-dependent acquisition of MS/MS data for the six most intense ions using stepped normalized collision energy of 20, 40, and 60 eV.

**Features annotations and statistical analysis of metabolomics datasets**

The raw data were processed with MS-DIAL version 4.70 for mass signal extraction between 100 and 1,500 Da from 0.5 to 10.6 minutes, respectively (Fraisier-Vannier et al 2020, Tsugawa et al 2015). MS1 and MS2 tolerance were set to 0.01 and 0.025 Da in the centroid mode. The optimized detection threshold was set to $5 \times 10^5$ concerning MS1 and 10 for MS2. Peaks were aligned on a QC reference file with a retention time tolerance of 0.15 minutes and a mass tolerance of 0.015 Da. Minimum peak height was set to 70% below the observed total ion chromatogram (TIC) baseline for a blank injection. Peak annotation was performed with an in-house database built on an MS-FINDER model (Fraisier-Vannier et al 2020).

MS-DIAL data were then cleaned with the MS-CleanR workflow by selecting all filters with a minimum blank ratio set to 0.8, a maximum relative standard deviation (RSD) set to 30, and a relative mass defect (RMD) ranging from 50 to 3.000. The maximum mass difference for feature relationships detection was set to 0.005 Da and the maximum RT difference to 0.025 min. Pearson correlation links were considered with correlation ≥ 0.8 and statistically significant with $\alpha = 0.05$. Two peaks were kept in each cluster*, viz.,* the most intense and the most connected. The kept features (m/z × RT pairs) were annotated with MS-FINDER version 3.52. The MS1 and MS2 tolerances were, respectively, set to 10 and 20 ppm. Formula finders were only processed with C, H, O, N, and S atoms. Databases (DBs) based on *Arabidopsis* (genus), *Brassicaceae* (family), and *Streptomyces* (genus of the AgN23 strain), were constituted with the dictionary of natural products (DNP, CRC press, DNP on DVD v. 28.2). The internal generic DBs from MS-FINDER used were KNApSAcK, PlantCyc, NANPDB, UNPD, COCONUT, and CheBI. Annotation prioritization was done by ranking *Arabidopsis* DB, followed by *Brassicaceae* DB, *Streptomyces* DB, and finally generic DBs using the final MS-CleanR step.

Statistical analyses were done by using SIMCA (version 14.1, Umerics). All data were scaled by pareto scaling before multivariate analysis. The (orthogonal) projection to latent structure using discriminant analysis ((O)PLS-DA) was used to separate data according to *A. thaliana* growing conditions. Variable Importance in Projection (VIP) features were selected according to their VIP score (>3). The principal component analysis was built with the web-interface MetaboAnalyst version 5.0 (http://www.metaboanalyst.ca)(Pang et al 2021). Significantly different metabolites and affiliated chemical classes with ClassyFire were selected using the criteria of $p < 0.05$ (t-test, control vs. treatment, unadjusted p-value) and log2 fold change (log2FC) > 0.8 or < -0.8(Djoumbou Feunang et al 2016).

### REFERENCE


Allen DG, Blinks JR, Prendergast FG (1977). Aequorin Luminescence: Relation of Light Emission to Calcium Concentration-A Calcium-Independent Component. *Science* **195:** 996-998.

Clough SJ, Bent AF (1998). Floral dip: a simplified method for Agrobacterium-mediated transformation of Arabidopsis thaliana. *Plant J* **16:** 735-743.

Djoumbou Feunang Y, Eisner R, Knox C, Chepelev L, Hastings J, Owen G *et al* (2016). ClassyFire: automated chemical classification with a comprehensive, computable taxonomy. *Journal of Cheminformatics* **8:** 61.

Fraisier-Vannier O, Chervin J, Cabanac G, Puech-Pages V, Fournier S, Durand V *et al* (2020). MS-CleanR: A feature-filtering approach to improve annotation rate in untargeted LC-MS based metabolomics. Analytical Chemistry, **92**: 9971-9981.

Gayrard D, Nicolle C, Veyssière M, Adam K, Martinez Y, Vandecasteele C *et al* (2023). Genome Sequence of the Streptomyces Strain AgN23 Revealed Expansion and Acquisition of Gene Repertoires Potentially Involved in Biocontrol Activity and Rhizosphere Colonization. *PhytoFrontiers™***:** PHYTOFR-11-22-0131-R.

Johansson ON, Nilsson AK, Gustavsson MB, Backhaus T, Andersson MX, Ellerström M (2015). A quick and robust method for quantification of the hypersensitive response in plants. *PeerJ* **3:** e1469.

Mithöfer A, Mazars C, Maffei ME (2009). Probing Spatio-temporal Intracellular Calcium Variations in Plants. In: Pfannschmidt T (ed). *Plant Signal Transduction: Methods and Protocols*. Humana Press: Totowa, NJ. pp 79-92.

Pang Z, Chong J, Zhou G, de Lima Morais DA, Chang L, Barrette M *et al* (2021). MetaboAnalyst 5.0: narrowing the gap between raw spectra and functional insights. *Nucleic Acids Research* **49:** W388-W396.

Pauly N, Knight MR, Thuleau P, Graziana A, Muto S, Ranjeva R *et al* (2001). The nucleus together with the cytosol generates patterns of specific cellular calcium signatures in tobacco suspension culture cells. *Cell Calcium* **30:** 413-421.

Tsugawa H, Cajka T, Kind T, Ma Y, Higgins B, Ikeda K *et al* (2015). MS-DIAL: data-independent MS/MS deconvolution for comprehensive metabolome analysis. *Nature Methods* **12:** 523-526.
