## Supplementary figures and images for "Root-associated *Streptomyces* produce galbonolides to modulate plant immunity and promote rhizosphere colonisation"

### Supplementary movie: EM-CCD observations of calcium waves in the nuclei of an aequorin-expressing Arabidopsis seedling.

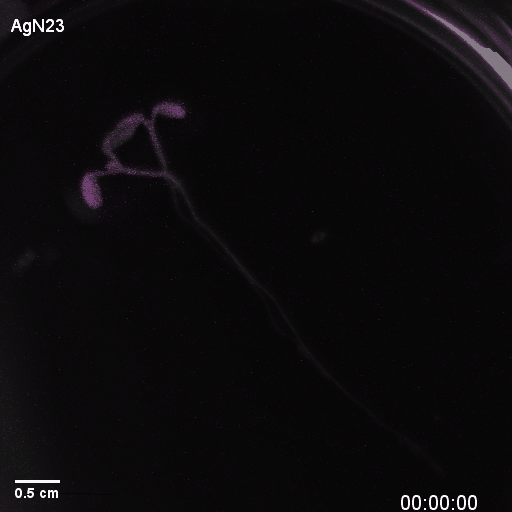
